# Constitutive, calcium-independent endoplasmic reticulum-plasma membrane contact site oscillations and its implications in store-operated calcium entry

**DOI:** 10.1101/2022.03.30.486443

**Authors:** Ding Xiong, Cheesan Tong, Yang Yang, Jeffery Yong, Min Wu

## Abstract

The endoplasmic reticulum and plasma membrane (ER-PM) contact site dynamics plays a central role for store-operated calcium entry (SOCE). ER localized calcium sensor STIM1 translocates to the contact sites, interacts with Orai and mediates calcium influx from the extracellular environment. Different species of phosphoinositides (PIPs) have been reported to be involved in contact site dynamics as well as STIM1 translocation. However, most of the studies were based on loss-of-function experiments or conditions that generate massive calcium store depletion. The kinetics of ER-PM contact site dynamics during physiological stimuli -induced calcium oscillations are not well understood. Using total internal reflection fluorescence microscopy (TIRFM), we investigated the relationship between dynamics of STIM1 as well as cortical ER (cER) proteins and calcium oscillations in rat basophilic leukemia (RBL) mast cells. Surprisingly, a significant percentage of cells displayed cyclic STIM1 and cER dynamics that were calcium-independent. Using specific lipid sensors, we showed that cyclic ER-PM contact site assembly was in phase with PI(4)P oscillation, but preceded phases of PI(4,5)P2 or PI(3,4,5)P3 oscillation. Optogenetic recruitment of the phosphoinositide 5-phosphatase from INPP5E, which decreased PI(4,5)P2 and increased PI(4)P levels on the plasma membrane, stimulated the translocation of STIM1 and inhibited calcium oscillations. Interestingly, prolonged stable translocation of STIM1 to the plasma membrane had an inhibitory effect on calcium oscillations. Collectively, our findings suggest that ER-PM contact sites formation is PI(4)P-dependent. In addition, reversibility of ER-PM contact sites dynamics and intermediate strength of ER-PM contact are needed for calcium oscillation.

## Introduction

Receptor activation of many cell types lead to biphasic calcium response with an initial phase of sustained calcium elevation, followed by oscillatory pulses(*1, 2*). Differentiating between the initial phase of sustained calcium elevation and calcium oscillations is necessary. Some cortical signaling modules, such as PKC (*3, 4*), F-BAR protein FBP17 and active Cdc42 (*5*), are coupled with calcium oscillations but not the initial sustained phase. In addition, there are frequency-modulated signaling and downstream consequences that are oscillation-specific (*2*). Hence, generalization using calcium as a universal second messenger is not sufficient and often misleading. Based on the waveforms of calcium oscillations, it has been recognized that there are at least two types of oscillatory responses (*6*). One is approximately sinusoidal and commonly superimposed on an elevated calcium level (*7, 8*). For the second type, calcium falls to resting levels for many seconds between spikes. It has been hypothesized that the sinusoidal oscillations are induced through a cytoplasmic oscillator involved in calcium-induced calcium release, while the second, baseline spiking type are associated with receptor activation and other events at the plasma membrane that mediate calcium entry (*6, 9, 10*). However, this classification is likely overly simplistic as baseline calcium spiking were also observed in the absence of external calcium (*8, 11*). Nevertheless, the functional consequences of calcium oscillations likely differ depending on whether SOCE is involved (*12*), which further impinges on the factors involved in specifically regulating SOCE.

A periodic activation of store-operated channels (SOC) by calcium store sensor STIM have been proposed to support a robust periodic calcium influx (*13*-*16*), but dynamics of STIM activity during calcium oscillations was much less characterized compared to other relatively non-physiological conditions such as thapsigargin-induced massive store depletion. Although it is widely believed that SOCE requires the negatively charged lipids, possibly phosphoinositides, it remains unclear whether the requirement is for STIM1 targeting to these contact sites or Orai1 gating and activation (*1*). There are mixed results regarding the specificity of the phosphoinositides required (*1*). The cytosolic portion of STIM1 contains a lysine-rich polybasic domain which is responsible for phospholipid interaction at the plasma membrane (*17*). *In vitro*, the polybasic segment of STIM1 binds both PI(4,5)P2 and PI(3,4,5)P3 with higher preference for PI(4,5)P2 than PI(3,4,5)P3 (*18*), but also has activity towards PI(3,4)P2, PI(3,5)P2 (*19*). It was also shown that a plasma membrane-like environment with lipids such as phosphatidylethanolamine, phosphatidylserine and phosphatidylinositol is required in addition to PI(4,5)P2 (*18*). Studies on the STIM1 translocation *in vivo* also show mixed results. PI(4,5)P2 and PI(3,4,5)P3 (*17*), PI(4)P (*20, 21*), or a combination of PI4P, PI(4,5)P2 and PI(3,4,5)P3 (*20*) have all been proposed to be the main determinant of SOCE. Besides binding to PI(4,5)P2, STIM1 is also proposed to transfer between PI(4,5)P2-rich and PI(4,5)P2-poor domains (*22*). These mixed results could be due to the fact that most of the current studies were based on loss-of-function experiments. The loss of SOCE upon artificial reduction of a given lipid indicates that this lipid at basal level contributed to SOCE but does not imply it is the rate-limiting step in the event of activation.

Here we aim to characterize the kinetics of cortical ER (cER) and STIM1 translocation in the context of receptor-mediated calcium oscillations, and to examine which plasma membrane cue defines the rate-limiting step during the periodic activation of STIM1/Orai1. Mast cells play vital roles in innate and adaptive immunity which are partially based on intracellular calcium signaling (*23*). Using RBL-2H3 mast cell, a well-established model system for calcium signaling (*24*), we show that the cyclic translocation pattern of STIM1 and cER exist in two forms. In particular, a periodic wave of STIM1 and cER translocation occurs without calcium oscillations. The oscillatory translocation of STIM1 is in sync with PI(4)P oscillations, but not PI(4,5)P2. We applied optogenetic tools to acutely increase PI(4)P and obtained gain-of-function evidence for its role in the kinetics of STIM1 translocation and calcium oscillations. Lastly, optogenetic clamping and stabilization of STIM1 showed a direct inhibitory effect on calcium oscillations, suggesting the importance of the reversibility of its translocation.

## Results

### Quantitative assessment of the requirement of SOCE in calcium oscillations

In secretory RBL-2H3 mast cells, multivalent antigen elicited a dramatic cytosolic Ca^2+^ increase followed with calcium oscillations (**Fig. 1a, Supplemental video 1**). To monitor the calcium oscillations, we established a RBL-2H3 cell line stably expressing genetically encoded calcium indicator GCaMP3. The population of cells with lowest fluorescence signal was selected using FACS to minimize effects due to sensor overexpression. Given the complexity of calcium oscillations and heterogeneity in the original source of calcium leading to cytoplasmic calcium elevation, it remains difficult to determine unambiguously *in situ*, whether a given cycle within a train of calcium oscillations rely on SOCE or not. We therefore set out to determine how to define quantitative criteria that facilitate the distinction between SOCE and alternative forms of calcium oscillations. As the intracellular stores have a relatively small capacity(*25*), the duration of calcium oscillations is likely short in the absence of SOCE (*24, 26, 27*). We confirmed the dependence of calcium oscillations on SOCE in cells pre-incubated in calcium-free buffer with 0.5mM EGTA chelator, a condition that reduced extracellular Ca^2+^ concentration to be about 10 nM. Antigen stimulation in the presence of EGTA could only elicited transient calcium oscillations, lasting an average of 2.9±1.2 cycles (148 cells from 6 experiments) (**Fig. 1b-c**), and is consistent with previous reports of RBL-2H3 cells (*24, 26*) or related RBL-1 cells (*12, 28*). When the calcium-free buffer was exchanged for one containing 1.8mM calcium, we observed robust long-lasting calcium oscillations in a majority of the cells (**Fig. 1b**). Consistent with our previous study (*5*), a biphasic response could be resolved after supplying extracellular calcium. The initial phase displayed an elevated baseline lasting between 3 to 7 min, while the second phase predominantly displayed baseline spiking type of calcium oscillations (**Fig. 1b**).

**Fig. 1.**
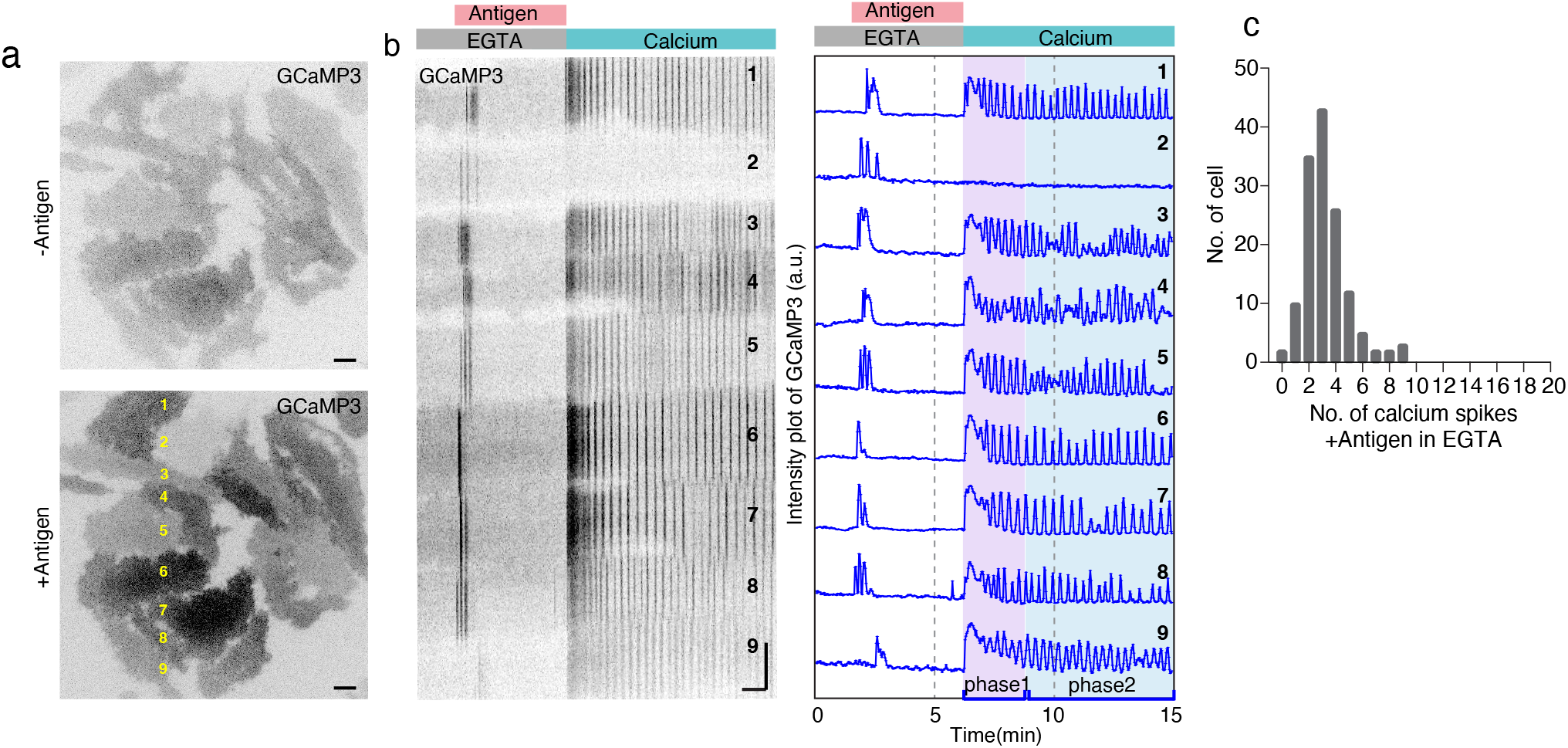
Determination of the upper limit of calcium pulses in the absence of extracellular calcium. (a) TIRF images of RBL-2H3 cells stably expressing GCaMP3 sensitized with anti-DNP IgE for overnight, and stimulated by antigen DNP-BSA in calcium-free buffer with 0.5mM EGTA and exchange to calcium-containing buffer (1.8mM Ca^2+^) during imaging. (b) Kymographs and time-series profiles of the cells marked in yellow in (a), showing calcium oscillations in the absence or presence of extracellular calcium. Phase 1 and phase 2 are marked respectively. (c) Histogram showing the number of calcium spikes upon DNP stimulation in the absence of extracellular calcium (148 cells from 6 experiments). Images are acquired 2sec/frame, grayscale images and kymographs are shown with inverted lookup table. For TIRF images, scale bar=10µm; for kymographs, scale bar=1min (horizontal bar), 10µm (vertical bar).

We also tested the effects of adding EGTA to cells displaying sustained calcium oscillation (**Supplemental Fig. 1a**). With initial stimulation with antigen, about 92% cells (174 out of 189 cells, 5 experiments) displayed calcium oscillations while the rest showed a single broad peak without calcium oscillations. Upon EGTA treatment, 23% of the cells (40 out of 174 cells) stopped their calcium oscillations completely, while 76% of the cells (133 out of 174 cells) displayed some residual calcium responses, lasting on average 3.4±3.3 cycles **(Supplemental Fig. 1b)**. We observed rarely (one in 174 cells) cell that displayed more than ten cycles of calcium oscillations in EGTA-containing buffer **(Supplemental Fig. 1a)**. The number of spikes of calcium oscillations in the presence of EGTA was not statistically different if the EGTA treatment was together or after stimulation. Most cells could recover and resume oscillations when calcium was restored (**Supplemental Fig. 1a**). Taken together, these data suggest that the capacity of the store is sufficient to last three or four cycles without refill, and that under physiological conditions, sustained calcium oscillations do not deplete the store.

### Cyclic STIM1 translocation can be coupled or uncoupled from calcium oscillations

STIM1 is the major regulator for supporting calcium oscillations operated by SOCE^12^. A critical step for SOCE activation is the translocation of STIM1 to ER-PM contact sites where it binds to and activates the Orai1 calcium channel. First, we visualized the localization of STIM1 using TIRFM in antigen-stimulated cells. The intensity of STIM1 showed a transient increase upon antigen stimulation which was followed by cyclic patterns (**Fig. 2a**). To study the relationship between STIM1 translocation and calcium oscillations, we imaged STIM1 together with GCaMP3. Interestingly, we found that STIM1 oscillations were not always coupled with calcium oscillations. We first classified our dataset based on whether calcium displayed sustained oscillations (defined as a train of continuous spikes of more than ten cycles) in order to better attribute them to SOCE. To classify whether any given trace has oscillations quantitatively, we visually inspected the traces and also applied Fourier transform to the intensity profiles, where presence of peaks in the Fourier space was used to confirm oscillations. We excluded data from the first ten minutes after stimulation in our analysis because calcium oscillations during this period of time usually occur with an elevated baseline level and display sinusoidal shape, which could be indicative of their intracellular origins **(Fig. 1b).** Based on the fact that calcium oscillations do not last more than ten cycles in EGTA **(Fig. 1c, Supplemental Fig. 1b)**, sustained oscillations is a strong indication for a requirement of SOCE, although it does not imply that every single spike in the train requires external calcium or that external calcium is the only source for a given spike. While it is difficult to determine unambiguously *in situ* whether a given pulse within the train of calcium oscillations depends on extracellular calcium, these criteria should largely eliminate calcium oscillations due to solely intracellular oscillations without any contribution from SOCE.

**Fig. 2.**
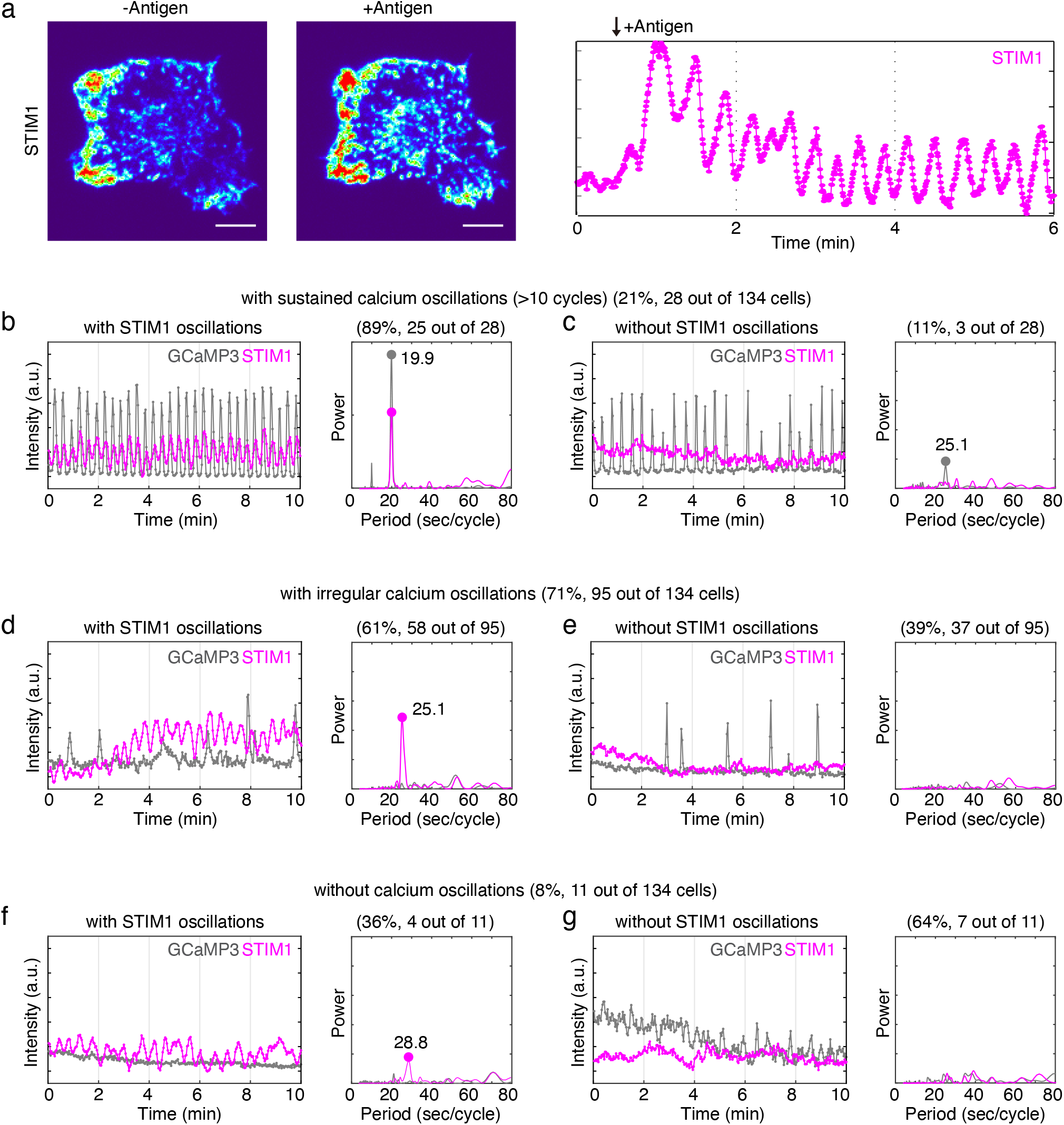
Cyclic STIM pattern is not necessarily coupled with calcium oscillations. (a) Cyclic STIM1 translocation upon antigen stimulation. *Left*: TIRF images of RBL-2H3 cell expressing STIM1-RFP before (-Antigen) and after (+Antigen) DNP-BSA stimulation. Images are shown with Thermal lookup table for improved clarity. *Right*: Representative STIM1 intensity profile of a cell being stimulated by antigen (n= 36 cells from 12 experiments). (b-g) Profiles of representative cells from various categories. Videos recording STIM1 and GCaMP3 signal simultaneously 2sec/frame for 10min (n=134 cells from 4 experiments). *Left*: Intensity profiles of STIM1-RFP and GCaMP3 fluorescence after antigen stimulation for 10-40min. *Right*: Cycling time of STIM1 and GCaMP3 oscillations as analyzed by fast-Fourier transform, with the periods of major eaks labelled. (b) Cell with robust STIM1 and calcium oscillations. (c) Cell with robust calcium oscillations without STIM1 oscillations. (d) Cell with irregular calcium oscillations but with STIM1 oscillations. (e) Cell with irregular calcium and no STIM1 oscillations. (f) Cell with no calcium oscillations but with STIM1 oscillations. (g) Cell showing no calcium or STIM1 oscillations. Scale bar=10µm.

Of the cells with robust, sustained calcium oscillations (21% of the cells, 28 out of 134 cells, 4 experiments), 89% (25 out of these 28 cells) showed coupled STIM1 oscillations (**Fig. 2b, Supplemental video 2**). The coupled STIM1 and calcium oscillations showed matching frequencies **(Fig. 2b)**, whereas cells with only calcium oscillations show a major peak for calcium but not STIM1 **(Fig. 2c)**. In 71% of cells (95 out of 134), calcium spikes were less regular (train of continuous oscillations < ten cycles). In these cells, 61% (58 out of 95) displayed STIM1 oscillations **(Fig. 2d-e, Supplemental video 2)**. There were also 8% (11/134) cells that did not have any calcium spike, and 36% (4/11) of them showed STIM1 oscillations (**Fig. 2f-g, Supplemental video 2)**. Consistent with the observation that STIM1 oscillations could take place in cells without calcium oscillations, we have also observed cyclic STIM1 patterns in cells stimulated in EGTA-containing buffer (**Supplemental Fig. 1c**). These data suggest that in cells with robust calcium oscillations where SOCE is relevant, STIM1 oscillations are prevalent **(Fig. 2b)**, but even in cells without calcium oscillations, STIM1 oscillations are common as well, indicating a possible calcium-independent, constitutive form of STIM1 dynamics **(Fig. 2d)**.

### Diverse patterns of cyclic STIM1 translocation

STIM1 translocation has been frequently studied in the context of store depletion using chemical inhibitors that block ER store refilling (namely thapsigargin and ionomycin), but has not been thoroughly investigated in physiological stimuli-induced calcium oscillations. Using calcium oscillations elicited by agonist stimulation of HEK293 cells as a model, STIM1 oscillations were observed in a small subset of the cells (10/91 cells) (*29*). Our results of the calcium oscillation-coupled STIM1 oscillations are therefore consistent with Bird et al., but the proportion of such couplings were higher in our experiments. In addition, STIM1 oscillations in cells without calcium oscillations or no calcium responses were unexpected. STIM1 functions as a calcium sensor in the ER and its translocation to cortical region has been assumed to be a result of store depletion. If STIM1 translocation to the plasma membrane can happen without changes in cytoplasmic calcium concentration, cyclic STIM1 translocation to the cortex may not be simply a calcium-dependent downstream response. Upon closer examinations of the cyclic STIM1 translocation patterns that were coupled with or without calcium oscillations, we found that the patterns of STIM1 dynamics are different under these two conditions. When there are calcium oscillations, STIM1 translocation takes place synchronously in different regions of the same cell and appeared as standing waves (**Fig. 3a, Supplemental video 2**). When calcium oscillations are absent, STIM1 puncta translocation was asynchronous in different regions across the cell, appearing as travelling waves (**Fig. 3b, Supplemental video 2**). Interconversion of these two patterns can be observed in cells where calcium oscillations became more intermittent while cyclic STIM1 translocation persisted **(Fig. 3c)**, suggesting that they are both physiological states of the same cell, rather than due to cell-to-cell variations.

**Fig. 3.**
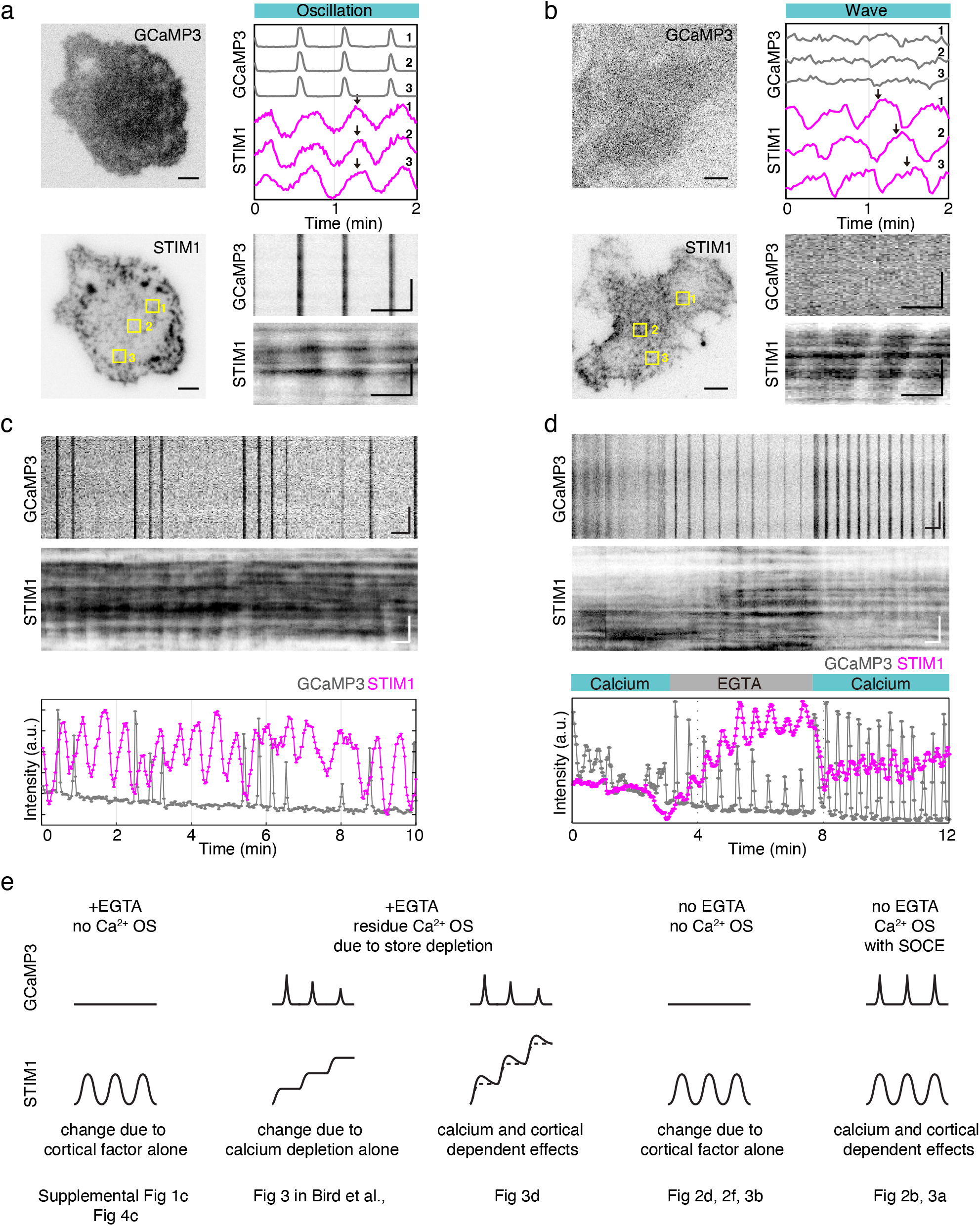
Diverse cyclic patterns of STIM1 translocation suggest regulation by other factors in addition to calcium. (a-b) Representative cells and profiles with STIM1 oscillations or waves, 15min after antigen stimulation. *Left*: TIRF images of a cell expressing STIM1-RFP and GCaMP3. *Top right*: Intensity profiles of GCaMP3 and STIM1-RFP fluorescence intensity at indicated regions on the left panel. Arrowheads showed the synchronized or temporally-shifted peaks of STIM1. *Bottom right*: Kymographs of GCaMP3 and STIM1-RFP from the same cell. For STIM1 oscillations, n= 16 cells; for STIM1 waves, n=7 cells. (c and d) Kymographs with intensity profiles of a cell expressing GCaMP3 and STIM1-RFP showing intermittent calcium oscillations with sustained STIM1 oscillations, 40min after antigen stimulation (c) (n=19 cells from 4 experiments) and a cell briefly starved of extracellular calcium (EGTA) (n=3 cells from 3 experiments), 10min after antigen stimulation (d). (e) Schematic illustration of the observed patterns in the profiles of STIM1 and calcium oscillations (Ca^2+^ OS) with the potential underlying causes. For TIRF images, scale bar=5 µm; for kymographs, scale bar=30 sec (horizontal bar), 5 µm (vertical bar).

The cyclic, calcium-independent STIM1 translocation we observed are likely separate phenomena from another interesting observation Bird et al., reported, which is that the translocation of STIM1 seemed to be amplified in the absence of extracellular calcium. To illustrate the distinction, we took advantage of the residual calcium oscillations (likely of intracellular origin) when stimulated cells were treated with EGTA. In the subset of cells with residual calcium oscillations, cyclic STIM1 translocation could be observed. Interestingly, as calcium oscillations gradually reduced in intensity, the baseline of cyclic STIM1 patterns increased **(Fig. 3d)**. These cells likely contain bigger calcium internal stores that could sustain a few pulses of calcium oscillations without refill, but with each pulse of cytoplasmic calcium, the ER store is incrementally depleted, leading to an increase of the STIM1 translocation to the plasma membrane, as reflected by the changes in the baseline level of the STIM1 pulses. These changes in baselines are likely reminiscent of the staircase-like increase of STIM1 translocation reported in Bird et al. using HEK293 cells. Here we found that STIM1 oscillations in the absence of SOCE are oscillations superimposed on top of changes of the rising baseline levels. They differed from STIM1 oscillations observed when calcium was reintroduced **(Fig. 3d)** or when calcium oscillations were not coupled **(Fig. 3b,c**). In the case of normal store refill, STIM1 likely returned to baseline after each pulse. Hence, this staircase-like increase in STIM1 baseline level could provide a signature to assess the extent of STIM1 translocation contributed by store depletion (**Fig. 3e**). However, there must be an additional mechanism regulating the reversible cortical translocation of STIM1 on top of the rising baselines.

### Cyclic STIM1 translocations are coupled with ER-PM contact site dynamics

The absence of cytoplasmic calcium changes argues for additional mechanisms underlying reversible STIM1 translocation to the cortex that are distinct from those regulating STIM1 translocation through store depletion. When STIM1 and calcium oscillations are coupled, it would be challenging to differentiate the effect of transient ER calcium store depletion from that of other cortical factors in STIM1 translocation. For instance, thapsigargin has been widely used to induce massive cortical STIM1 accumulation, but thapsigargin also causes the maximal depletion of ER stores, with the resulting influx of calcium into the cytosol, and impact on STIM1 oligomerization and phosphorylation, could overwhelm any other cortical effects. We therefore used STIM1 waves as a system to pinpoint the effect of cortical factors on STIM1 translocation from that of calcium store depletion.

As STIM1 is a transmembrane ER protein, we visualized the dynamics of STIM1 together with ER marker ER5a to determine whether similar dynamics happened to other ER protein at the cell cortex. Upon antigen stimulation, we observed cyclic translocation of ER5a in the TIRF field that were coupled with STIM1 (**Fig. 4a, Supplemental video 3**). Temporal profile of the TIRF images showed similar level of clustering for STIM1 and ER5a oscillations (**Fig. 4a**), which is in contrast to the well-established pattern of STIM1 enrichment when calcium store was depleted by thapsigargin (**Fig. 4b**). Another ER marker Sec61 also displayed waves when it was expressed alone and their cyclic translocation to the cortical region persisted with similar amplitude when extracellular calcium was depleted by EGTA treatment, further supporting the occurrence of constitutive STIM1 and cER oscillations in a calcium-independent manner (**Fig. 4c**). ER-PM contact marker MAPPER (*30*) also displayed similar oscillations, confirming dynamic ER-PM contact sites formation (**Fig. 4d**). These results suggest that the cyclic STIM1 translocation reflect dynamic ER-PM contact site dynamics that did not show a selective enrichment of STIM1 relative to other ER proteins such as ER5a and Sec61, likely because there was no significant store depletion under such condition. Although under severe condition of store depletion STIM1 could be selectively enriched at ER-PM junction, constitutive oscillations of cortical ER are sufficient for STIM1 translocation during physiological stimulations.

**Fig. 4.**
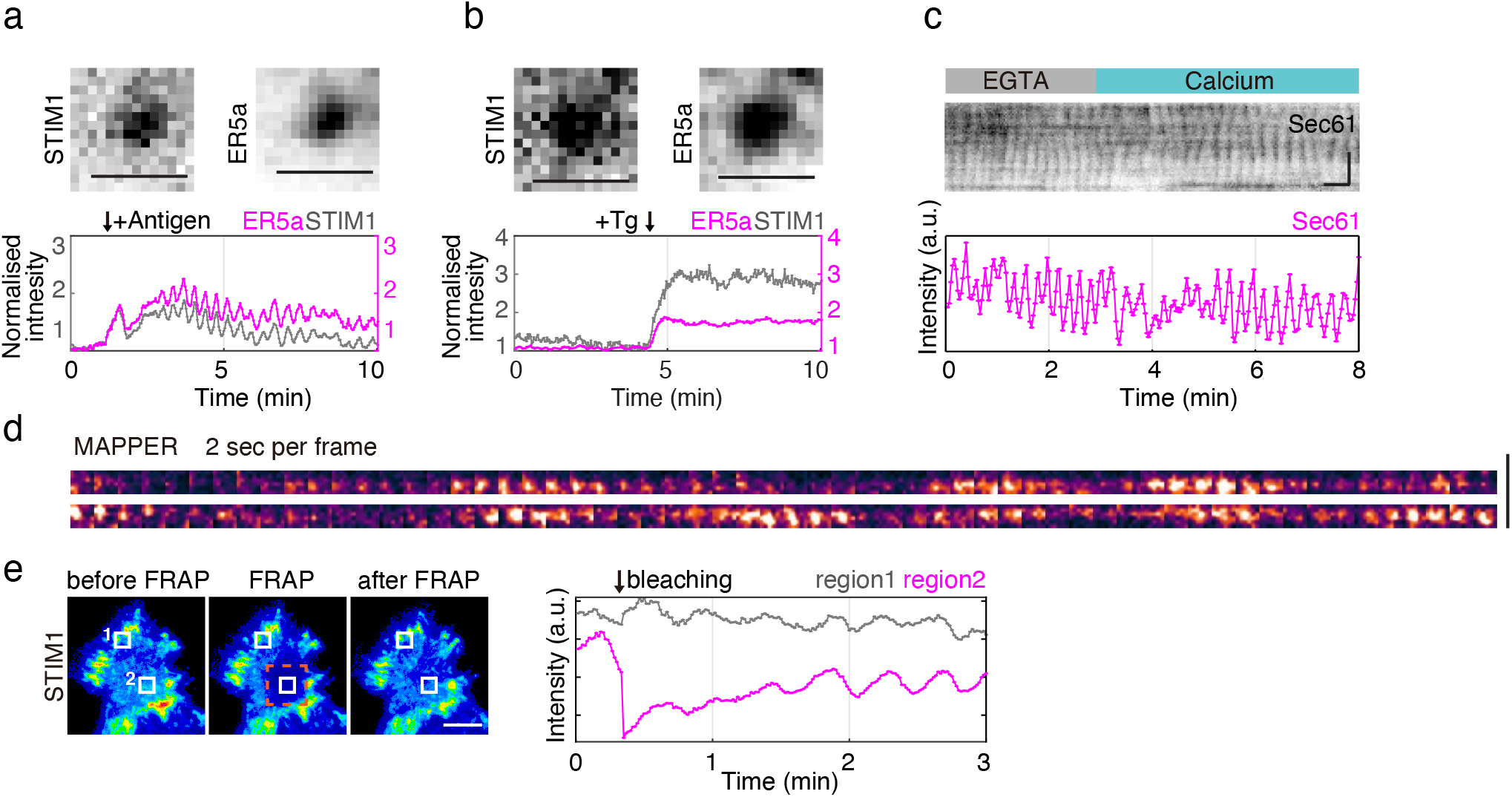
Cyclic STIM1 translocations are synchronized with cortical ER-PM contact site dynamics. (a) TIRF images (*upper*) and intensity profile (*bottom*) of mEGFP-ER-5a and STIM1-RFP individual puncta in wave upon antigen stimulation (n= 5 cells from 4 experiments). (b) TIRF images (*upper*) and intensity profile (*bottom*) of STIM1-RFP and mEGFP-ER-5a individual puncta in wave upon 2*μ*M thapsigargin (n=8 cells from 4 experiments). (c) Representative kymograph and intensity profile of mCherry-Sec61 showing ER waves in the absence or presence of extracellular calcium, 10min after antigen stimulation (n=5 cells from 2 experiments). (d) Montage of 60 frames (2sec interval) showing oscillatory dynamics of individual GFP-MAPPER puncta after antigen stimulation (n=9 cells from 3 experiments). (e) Fluorescence recovery after photobleaching (FRAP) of STIM1-RFP. Pseudocolored TIRF images before and after photobleaching (red square) with the intensities of two regions boxed in white were analyzed. The experiment was carried out 30min after antigen stimulation (25 cells from 3 experiments). For TIRF images and montages, scale bar= 5 µm; for single puncta images in Fig. 4a-b, scale bar=1 µm; for kymographs, scale bar=30 sec (horizontal bar), 5 µm (vertical bar).

We then studied the dynamics of STIM1 puncta using fluorescence recovery after photobleaching (FRAP) (**Fig. 4e**). Upon bleaching, STIM1 signal largely recovered, with a relatively small immobile fraction (13 ± 8%), suggesting that the accumulation in the cortical region was due to preferential clustering that is dynamically maintained. The kinetics of recovery (45 ± 20sec) was slower than the oscillation cycle of STIM1 (22 ± 3sec), further suggesting that oscillations were regulated by separate kinetic events from lateral diffusion within the ER membrane. We also tested whether STIM1 oscillations required its polybasic domain, which is thought to be important for its plasma membrane targeting. We found STIM1-Δpolybasic domain displayed a decreased oscillation amplitude (52±14%) compared to STIM1-RFP if Orai1 was overexpressed (**Supplemental Fig. 2a**), but similar oscillation amplitude without Orai1 overexpression (**Supplemental Fig. 2b**). Regardless of whether Orai1 was overexpressed, the percentage of cells displaying STIM1-Δpolybasic domain oscillations was reduced relative to that of STIM1-RFP (**Supplemental Fig 2c, d,** from 44±10% to 21±8% without Orai1 overexpression, from 54±4% to 37±5% with Orai1 overexpression).

### PI(4,5)P2 is not rate-limiting for STIM1 and ER-PM contact site oscillations

Previous *in vitro* studies showed that STIM1 binds to PI(4,5)P2 and PI(3,4,5)P3 (*18, 19*). To better understand the cortical translocation mechanisms of STIM1 and ER, we compared cyclic STIM1 translocation with other cortical events that depend on PI(4,5)P2. Synchronized oscillations and travelling waves of STIM1 are reminiscent of the waves of cortical actin, active Cdc42 and F-BAR protein FBP17 that we previously reported (*5*). We had shown that recruitment of FBP17 and N-WASP are dependent on the *de novo* synthesized PI(4,5)P2, which cycles with similar phases as FBP17 and N-WASP (*31*). We therefore investigated the relationship between STIM1 and FBP17/N-WASP. We found that STIM1 waves were coupled with FBP17 waves (**Fig. 5a, Supplemental video 4**). They shared the same periodicity, but peaks of STIM1 were about 4 sec prior to that of FBP17 (**Fig. 5a**). Similar temporal correlation exists between MAPPER and FBP17 (**Fig. 5b**), and between STIM1 and N-WASP (**Supplemental Fig. 3**). We also examined the dynamics of Septin4 which is essential for SOCE (*32*). Septin4 appeared in the waves, and was peaked about 6 sec prior to the corresponding FBP17 puncta (**Fig. 5c**). These phase-shifted cycles of translocation suggest that these dynamics were not due to random global membrane fluctuations. Cycles of STIM1, MAPPER and Septin4 translocation preceded that of PI(4,5)P2 effectors FBP17 and N-WASP, indicating that constitutive STIM1 and cER translocation appeared prior to the rise of PI(4,5)P2.

**Fig. 5.**
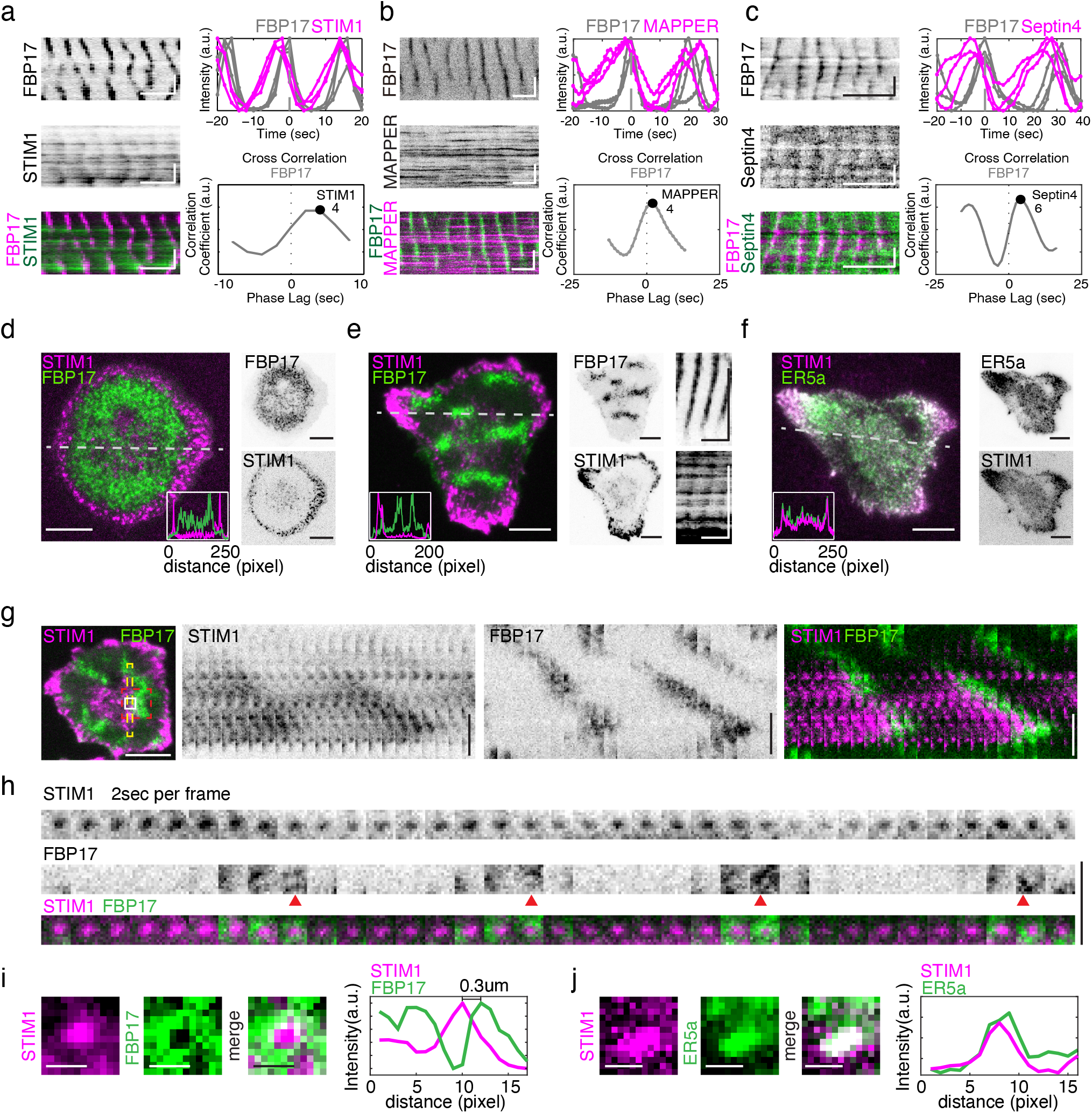
Coupling of ER proteins with cortical proteins in wave dynamics suggests PI(4,5)P2 is not rate-limiting for STIM1 and ER translocation. Kymographs, intensity profile of aligned peaks and cross-correlation analysis for cells stimulated with antigen between 10-30min was generated from a movie for (a) FBP17-EGFP and STIM-RFP (n= 20 cells from 5 experiments); (b) GFP-MAPPER and FBP17-EGFP waves (3 cells from 3 experiments); (c) mCherry-FBP17 and EGFP-Septin4 (n=12 cells from 4 experiments). (d-e) TIRF images of FBP17-EGFP (green) and STIM1-RFP (magenta) from representative cells without (d) or with (e) waves, 15min after antigen stimulation (for cell without waves, n=4 cells, for cell with waves, n=9 cells). Insets are line intensity profiles along the white dashed lines. (f) TIRF images of ER5a-GFP and STIM1-RFP (magenta) from representative cells 10min after antigen stimulation (5 cells from 4 experiments). Insets are line intensity profiles along the white dashed lines. (g) Representative image and montages of a cell displaying FBP17-EGFP (green) and STIM1-RFP (magenta) waves, 15min after antigen stimulation. *Left*: TIRF image. *Right*: Montages of 22 frames (2sec interval) of a ROI marked by yellow rectangle. (n= 20 cells from 5 experiments). (h) Montages for a single punctum of 35 frames (2sec interval) of region marked by white square in (g). Red arrows highlight the frames FBP17 in vicinity of STIM1 punctum. (i-j) Maximum projection view of 10 frames showing FBP17 surrounding STIM1 punctum, STIM1 colocalized with ER5a punctum. For TIRF images, scale bar=10 µm; for averaged puncta images in Fig. 5i-j, scale bar=1 µm; for montages in Fig. 5h, scale bar=5 µm; for kymographs, scale bar=1 min (horizontal bar), 5 µm (vertical bar).

There were additional distinctions between STIM1 and FBP17 patterns, even though waves of STIM1 and FBP17 were temporally and locally coupled. From the snapshots of cells co-expressing both markers, localization of STIM1 and FBP17 appeared to be almost mutually exclusive at a given time (**Fig. 5d-e**). Such exclusions were the most clear in cells without waves (likely cells with high PI(3,4,5)P3 according to our previous work (*31*)), where relatively stable FBP17 signals accumulated towards the centre of the cell, while STIM1 preferred the periphery (**Fig. 5d**). In a majority of cells, STIM1 was not restricted to the periphery but was still more concentrated around the edges, similar to ER5a (**Fig. 5e, f**). Furthermore, puncta of STIM1 and FBP17 that made up the oscillating waves did not register in space (**Fig. 5g)**. Close examination of oscillating STIM1 puncta revealed that FBP17 signals were recruited in the vicinity of STIM1 puncta, occupying distinct domains surrounding STIM1 (**Fig. 5h, i)**. In contrast, STIM1 and ER5a signals overlapped completely (**Fig. 5j).**

The opposite trend of amplitude fluctuations between FBP17 and STIM1 waves could also be seen right after stimulation, when FBP17 levels decreased while STIM1 increased in the same cell (**Supplemental Fig. 4a**). The intensity of STIM1 increased dramatically upon antigen stimulation (**Fig. 2a**), but PI(4,5)P2 (monitored by PH_PLC_) intensity dropped within the same period (**Supplemental Fig. 4b**). The translocation of STIM1 in spite of the acute, receptor-mediated PI(4,5)P2 hydrolysis is typically interpreted as due to changes in ER calcium concentrations, but it is also consistent with the possibility that PI(4,5)P2 levels may not be the rate-limiting step for STIM1 translocation.

The phase-shifted translocation of STIM1 and FBP17 proteins (**Fig. 5a**), their distinct localization pattern at the cellular level (**Fig. 5d-e**), and exclusive localization at the level of individual punctum (**Fig. 5h-i**) indicate to us that PI(4,5)P2 is neither rate-limiting for STIM1 puncta translocation, nor does it define the maximal amplitude of STIM1 translocation to the cortical region under our experimental conditions. This does not mean that PI(4,5)P2 is not necessary for STIM1 translocation. Rather, basal or even reduced levels of PI(4,5)P2 is sufficient for STIM1 translocation that supports calcium oscillations.

### Cyclic STIM1 translocations in sync with oscillations of PI(4)P but precede those of PI(4,5)P2 or PI(3,4,5)P3

PI(4)P has emerged as an important determinant of plasma membrane identity in addition to its role as a PI(4,5)P2 precursor (*21, 33*). To examine whether PI(4)P is the key phosphoinositide responsible for STIM1 translocation at the ER-PM contact sites, we imaged PI(4)P (monitored by tandem PH_Fapp1_) with STIM1. Interestingly, PI(4)P oscillated with the same phase as STIM1 whereas cytoplasmic protein iRFP, as a control for membrane fluctuation, did not show oscillations with either PI(4)P or STIM1 (**Fig. 6a**). When STIM1 cycle was compared against other phosphoinositide sensors, we observed that STIM1 was recruited prior to the rise of PI(4,5)P2 (monitored by PH_PLC_) (**Fig. 6b**), and PI(3,4,5)P3 (monitored by PH_Grp1_) (**Fig. 6c**). These are consistent with the relative phases of STIM1 compared with FBP17 and N-WASP (**Fig. 4a, Supplemental Fig. 3**) as both FBP17 and N-WASP shared similar phases with PI(4,5)P2 and PI(3,4,5)P3 (*31*). Tks4, a PI(3,4)P2-specific binding protein, displayed negative-correlation with STIM1 (**Fig. 6d**). These temporal relationships between STIM1 and phosphoinositides are consistent with PI(4)P levels being rate-limiting for STIM1 and ER-PM contact site oscillations (**Fig. 6e**).

**Fig. 6.**
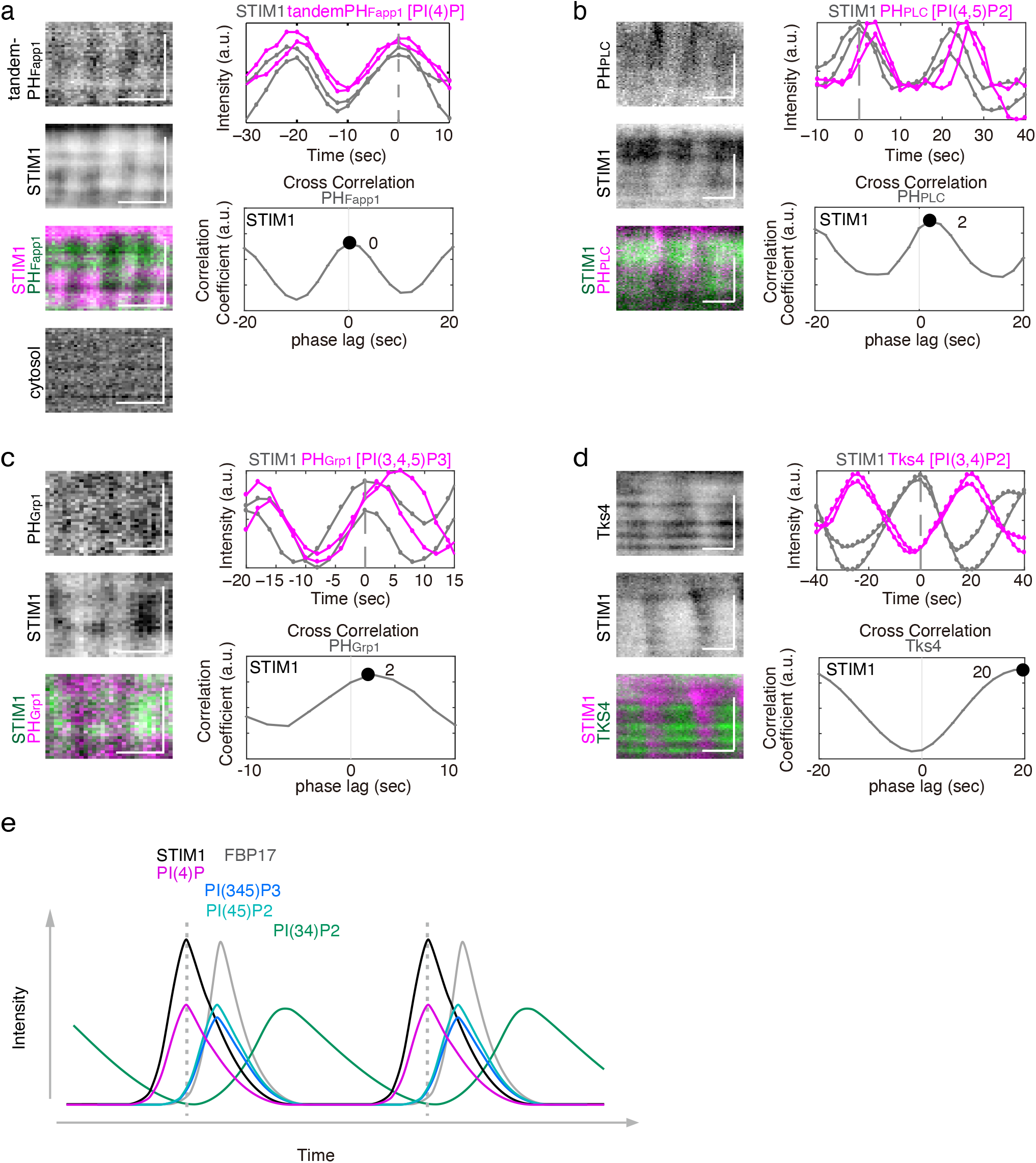
Oscillations of PI(4)P, but not PI(4,5)P2 or PI(3,4,5)P3 levels share the same phase with STIM1. (a-d) Kymographs, intensity profile and cross-correlation analysis of cells stimulated with antigen between 10-40mins for (a) STIM1-RFP and PI(4)P waves (monitored with EGFP-tandem PH_Fapp1_), iRFP as a Cytosol marker (n=9 cells from 4 experiments), (b) STIM1-RFP and PI(4,5)P2 (monitored with iRFP-PH_PLCδ_) (n=7 cells from 3 experiments), (c) STIM1-RFP and PI(3,4,5)P3 (monitored with mCherry-PH_Grp1_) (n=3 cells from 3 experiments), (d) STIM1-RFP and PI(3,4)P2 binding protein Tks4-GFP (n=11 cells from 3 experiments). (e) Schematic representation of the relationships between the oscillations of STIM1, FBP17 and the various phosphoinositides. STIM1 translocation is coupled with PI(4)P, followed by a delayed rise of PI(3,4,5)P3, PI(4,5)P2 and FBP17. PI(3,4)P2 is the last to rise and shares an reciprocal relationship with STIM1. Scale bar=30 sec (horizontal bar), 5 µm (vertical bar).

In order to test the hypothesis that PI(4)P, and not PI(4,5)P2, is the rate-limiting step of STIM1 translocation, we triggered the increase of PI(4)P and examined its effect on the kinetics of STIM1 translocation using optogenetics. We used genetically encoded light-inducible cryptochrome2 (Cry2) and plasma membrane targeted N-terminal portion of CIB1 (CIBN) dimerization to recruit the phosphoinositide 5-phosphatase domain of INPP5E (5-ptase_INPP5E_) to the plasma membrane (*34*), which should increase the level of PI(4)P while reducing the level of PI(4,5)P2 (**Fig. 7a**). Using this method, we could differentiate the roles of PI(4)P and PI(4,5)P2 on STIM1 and calcium oscillations. To confirm the changes in PI(4)P levels, we monitored its level with a specific marker P4M_SidM_. Upon blue light activation, dual-color recording showed that the plasma membrane recruitment of 5-ptase_INPP5E_ led to increased PI(4)P levels (**Fig. 7b**). By comparing the fluorescence intensity of P4M_SidM_ before and after blue light, the degree of increase in the PI(4)P intensity was about 33 ± 9%, while decreasing the intensity of PI(4,5)P2 (**Fig. 7c**), consistent with previous study (*31, 34*). Next, we examined the change of STIM1 upon the 5-ptase_INPP5E_ recruitment. The fluorescence intensity of STIM1 consistently increased after blue light activation (29 ± 6%) (**Fig. 7d**). The same strategy leads to reduction of FBP17 and N-WASP oscillation amplitude, both of which binds to PI(4,5)P2 (*31*). We also tested the effect of 5-ptase_INPP5E_ recruitment on calcium oscillations using genetically encoded calcium indicator GECO1. Increasing 5-ptase_INPP5E_ on the plasma membrane caused significant inhibition (**Fig. 7e, Supplemental video 5**) or complete abolishment (**Fig. 7f**) of calcium oscillations in 69% cells (9 of 13 cells, 4 experiments). The amplitude of spikes in calcium oscillations was significantly reduced by 46 ± 7% (95 and 90 peaks, respectively) (**Fig. 7e**). These results confirmed the known importance of PI(4,5)P2 in calcium oscillations (*35*) but suggests that its effect is not due to the lack of STIM1 translocation. The acute manipulation of PI(4)P levels and the corresponding decrease of PI(4,5)P2 strongly suggests that PI(4)P, not PI(4,5)P2, determines the kinetics of STIM1 translocation to the cortex.

**Fig. 7.**
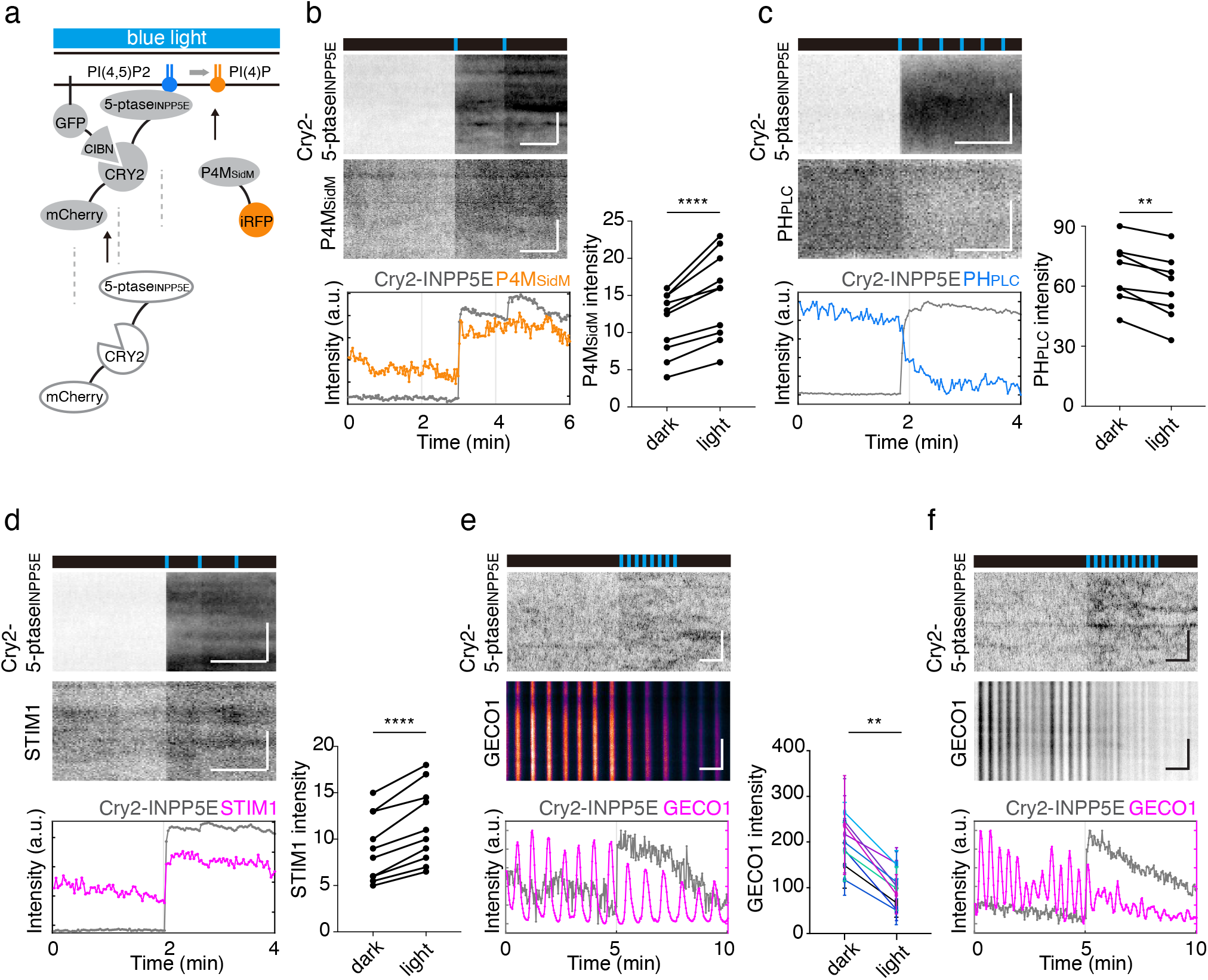
Modulation of STIM1 translocation and calcium oscillations by PI(4)P (a) Schematic of optogenetic light-induced plasma membrane recruitment of 5-phosphatase domain of INPP5E to generate PI(4)P. (b-d) *Left:* Representative kymographs and profiles before and after blue light pulses (2×114ms) for mCherry-CRY2-5-ptase_INPP5E_ and PI(4)P (monitored with iRFP-P4M_SidM_) (b), PI(4,5)P2 (monitored with iRFP-PHP_LCδ_) with 6×114ms pulses (c), STIM1-iRFP with 3×114ms pulses (d). *Right:* Quantification of lipid sensor or protein fluorescence intensity before and after blue light exposure for iRFP-P4M_SidM_ (n=12 cells from 3 experiments) (b), iRFP-PH_PLCδ_ (n=8 cells from 3 experiments) (c) and STIM1-iRFP (n=11 cells from 5 experiments) (d). (e-f) Representative kymographs and intensity profiles miRFP670-Cry2-ptase_INPP5E_ recruitment upon blue light leads to decreased amplitude with 8×114ms pulses (e) or inhibition with 10×114ms pulses (f) of calcium oscillations, 15min after antigen stimulation (n=9 cells from 4 experiments). Color plots showing the quantification for intensity of each spike in calcium oscillations before and after light pulses in different cells. (n= 95, 90 peaks, respectively). Pseudocolor is used for kymograph of GECO1 in (e) to emphasize the amplitude change. Error bar, s.d.; \*\**P*< 0.01, \*\*\*\**P*<0.0001, *t*-test. Scale bar=1 min (horizontal bar), 5 µm (vertical bar).

### Network hierarchy revealed by long-term oscillations

The uncoupling of cortical STIM1 translocation from calcium oscillations during optogenetic manipulation **(Fig. 7d-f)** as well as after antigen stimulation **(Supplemental Fig. 4a)** led us to hypothesize that STIM1 translocation and STIM1 activation of Orai1 are separate regulatory steps for SOCE, regulated by PI(4)P and PI(4,5)P2, respectively. If PI(4)P is a precursor of PI(4,5)P2, what is the benefit of such separation, especially if they are both cycling with the same periodicity? We found that while FBP17 and STIM1 waves were coupled on the short time scale of one oscillation cycle (20 seconds), amplitudes of these oscillations could fluctuate over a slower time scale (minutes) in different ways. Spontaneous changes in the amplitude of FBP17 and STIM1 were apparent when imaged over a duration of 20-30 min (**Fig. 8a)**. Their maximal intensities were often reciprocal. Increase of maximal STIM1 amplitude often correlated with decrease of maximal FBP17 amplitude over longer time scale while still retaining their relative phases within the shorter time scale (**Fig. 8a)**. In contrast, amplitude of FBP17 waves consistently correlated with active Cdc42 for both short and long time scale (**Fig. 8b, c)**. Tight coupling of FBP17 and active Cdc42 or PI(4,5)P2 oscillations on both short and long time scale is consistent with the idea that FBP17 is an effector of Cdc42 or PI(4,5)P2, and that FBP17 wave amplitude could be used as a readout for PI(4,5)P2 levels (*31*). Thus, the short time scale coupling between STIM1 and FBP17 (within each cycle) is likely consistent with the tight metabolic coupling of PI(4)P and PI(4,5)P2, while the opposing trendlines of STIM1 and FBP17 oscillations imply an additional, antagonistic mechanism occurring over longer time scale that uncouples the two **(Fig. 8d-e)**.

**Fig. 8.**
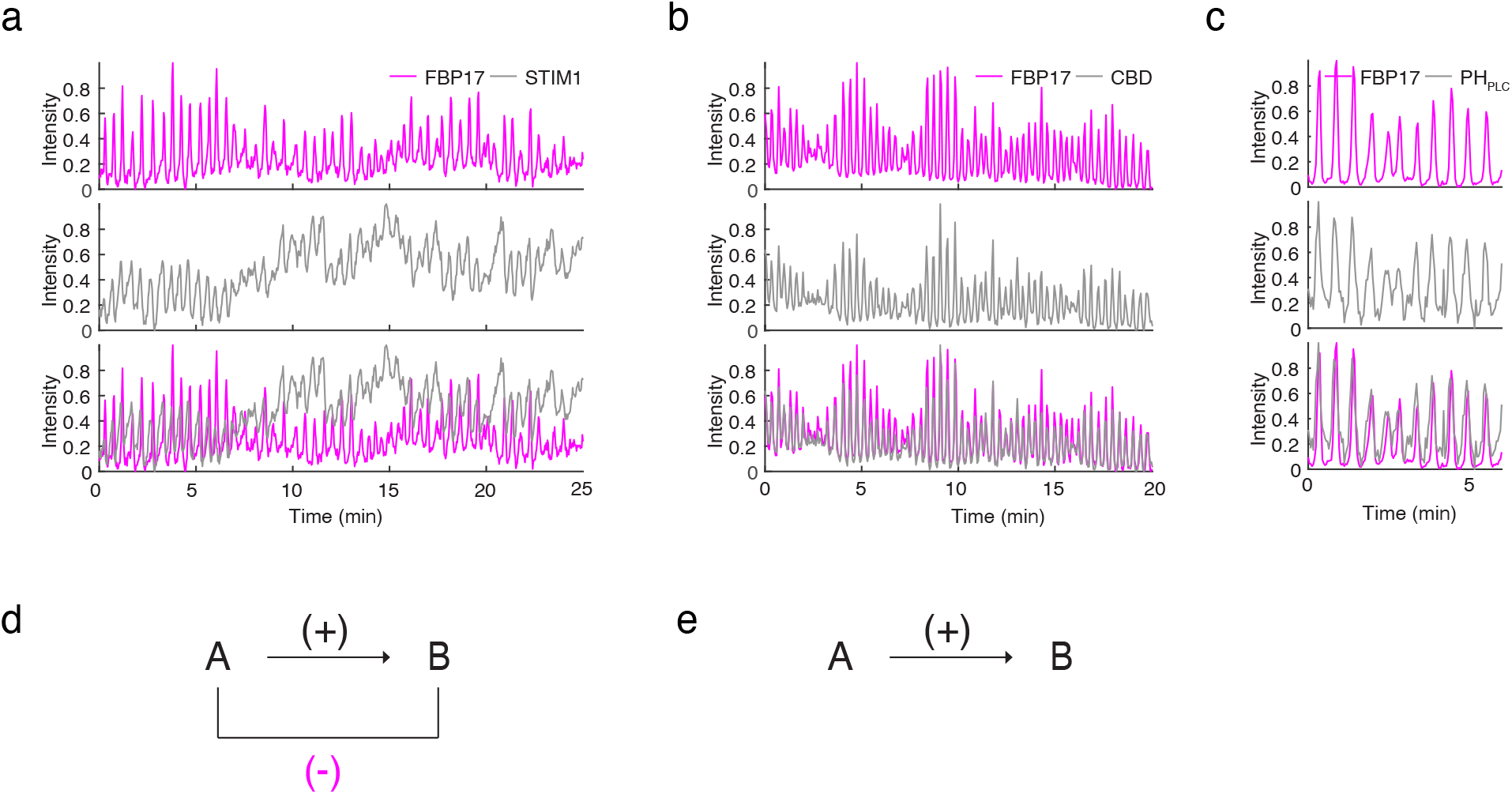
Network hierarchy revealed by long-term oscillations. (a-b) Intensity profiles of spontaneous changes of RBL-2H3 cells stably expressing FBP17-EGFP co-transfected with STIM1-RFP (a) (n=4 cells from 3 experiments), mRFP-wGBD (b) (n=8 cells from 3 experiments) oscillations within a 20-25min period, beginning after 10min of antigen stimulation. (c) Intensity profiles of spontaneous changes of cells stably expressing FBP17-EGFP co-transfected with iRFP-PH_PLCδ_ 10min after antigen stimulation (9 cells from 5 experiments). (d) Schematic illustration of the antagonistic relationship between of FBP17and STIM1. (e) Schematic illustration of the positive relationship between of FBP17 and active cdc42.

To test for the possible antagonistic regulation exerted by STIM1/ER-PM contact sites on calcium oscillations, we designed an optogenetic system to artificially clamp STIM1 to the cortex and stabilize it there by tagging STIM1 with Cry2 **(Fig. 9a)**. In addition to a 7 amino acid linker (GILQSTM), the long unstructured stretch of 243 amino acids after the SOAR domain of STIM1 would ensure flexibility for membrane binding after light induced translocation. We then tested the translocation of STIM1-Cry2-miRFP670 in cells transfected with plasma membrane localized CIBN using K-Ras derived CAAX motif (CIBN-GFP-CAAX). Upon blue light activation, STIM1-Cry2-miRFP670 was immediately translocated to the TIRF field as distinct puncta (**Fig. 9a**). To test the functionality of STIM1-Cry2-miRFP670, we examined its colocalization with STIM1-RFP. After blue light activation, we observed colocalization of STIM1-RFP with STIM1-Cry2-miRFP670 puncta, suggesting STIM1-Cry2-miRFP670 can also oligomerize with STIM1-RFP and bring them to ER-PM contact sites under light-induced translocation **(Fig. 9b, Supplemental video 6)**. The majority of STIM1 puncta existed prior to light-induction and increased in intensity after STIM1-Cry2-miRFP670 translocation, and a few were newly formed puncta **(Fig. 9b)**. In addition, there was a clear increase (40 ± 20%) of STIM1-RFP intensity together with STIM1-Cry2, indicating that the anchoring of STIM1-Cry2 to the membrane did not prevent other proteins from relocating there. We also tested the functionality of STIM1-Cry2 by examining its translocation to the plasma membrane upon thapsigargin treatment. STIM1-Cry2 formed clear puncta when cells were treated with thapsigargin without light induction, and had a significant intensity increase (115 ± 95%) **(Fig. 9c, Supplemental video 7)**.

**Fig. 9.**
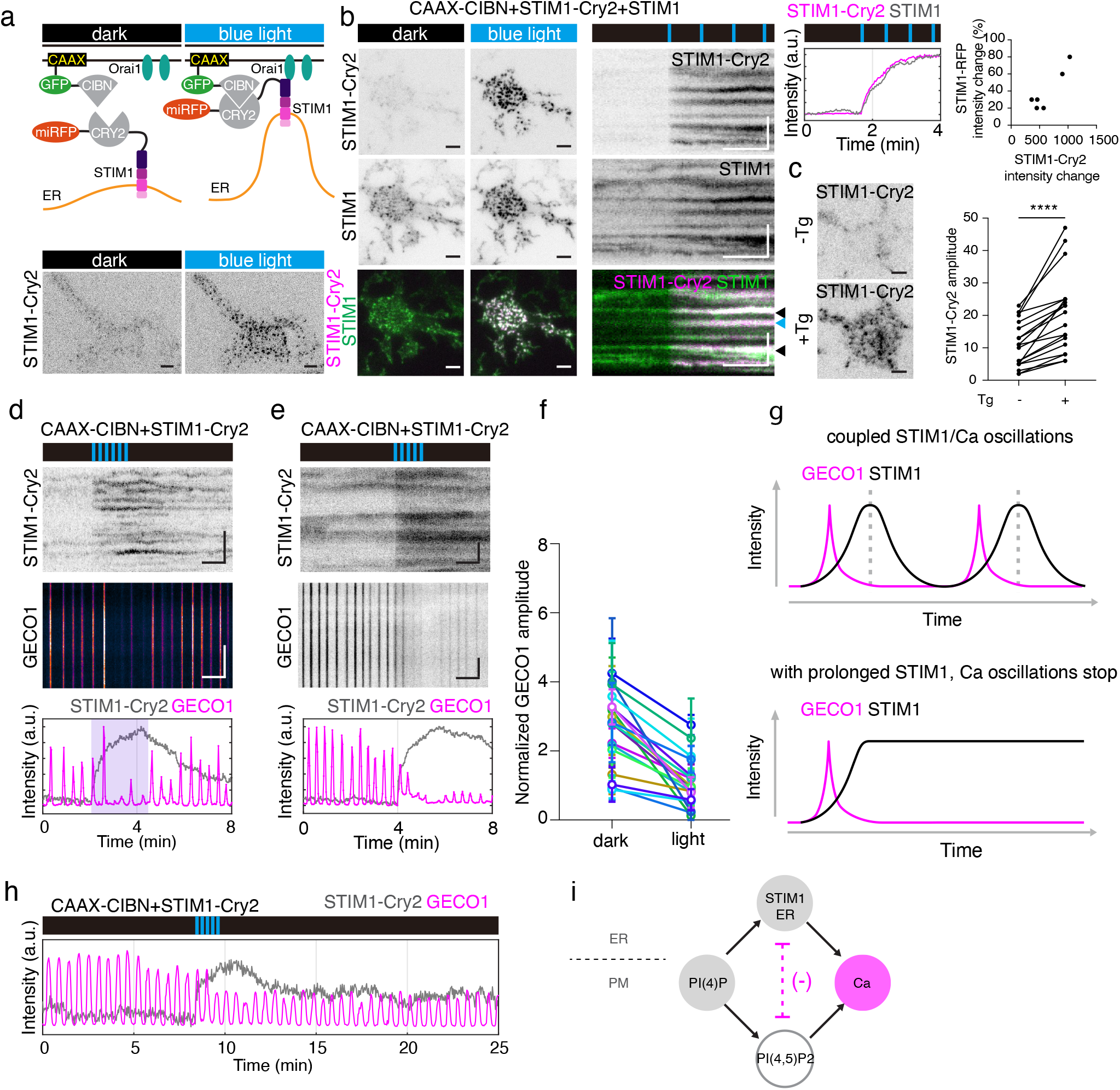
Optogenetic recruitment of STIM1 regulates calcium oscillations. (a) *Top*: Schematic of light-induced plasma membrane translocation of STIM1-Cry2-miRFP670 by CIBN-GFP-CAAX. *Bottom*: Representative TIRF images of a cell before and after blue light exposure without antigen stimulation (24 cells from 9 experiments). (b) TIRF images (*left*), kymographs (*middle*), intensity profile and scatter plot (*right*) showing coupled translocation of STIM1-Cry2-miRFP670 and STIM1-RFP upon blue light exposure, without antigen stimulation (n=6 cells). Black and blue arrows show pre-existing or newly formed STIM1 puncta, respectively. (c) TIRFM images (*left*) and quantification of STIM1-Cry2-miRFP670 intensity upon 10*μ*M thapsigargin (Tg) treatment, without antigen stimulation (19 cells from 3 experiments). (d-e) Kymographs and intensity profiles of STIM1-Cry2-miRFP670 and GECO1 during exposure to blue light pulses, movies were acquired 10 to 15min after antigen stimulation. Cell showing decreased (d) or almost abolished (e) calcium oscillations when STIM1-Cry2-miRFP670 is recruited upon blue light pulses. (f) Color plots showing the quantification for intensity of each spike in calcium oscillations before and after light pulses in different cells (n= 175, 167 peaks, respectively, 18 cells from 9 experiments). (g) Schematic illustration of the relationship between of STIM1 translocation and calcium oscillations. (h) Intensity profile of STIM1-Cry2-miRFP670 and GECO1 showed in a rare cell, robust but reduced amplitude of calcium oscillations with STIM1-Cry2 clamping. (i) Schematic illustration of the relationship between of STIM1/ER, calcium oscillations, PI(4)P and PI(4,5)P2. PI(4)P is a rate-limiting phospholipid for STIM1/ER plasma membrane cyclic translocation which could contribute to calcium oscillations. PI(4)P is also the substrate for PI(4,5)P2 synthesis whose hydrolysis leads to calcium oscillations. Increased recruitment of STIM1/ER could have an antagonistic effect on calcium oscillations. Pseudocolor is used for kymograph of GECO1 in Fig. 9d to emphasize the amplitude change. Error bar, s.d., **** *P*<0.0001, *t*-test. For TIRF images, scale bar=5 µm; for kymographs, scale bar=1min (horizontal bar), 5 µm (vertical bar).

Next, we examined the effect of stably clamping STIM1 on calcium oscillations. We found that sustained calcium oscillations were decreased or inhibited in 69% of cells (18 of 26 cells, 9 experiments). 15 of 18 cells showed decreased maximal GECO1 intensity (**Fig. 9d**), while 3 of 18 cells showed near complete inhibition of calcium oscillations (**Fig. 9e**). The decreased amplitude could recover as blue light pulses were removed (**Fig. 9d**). The degree of decrease in the amplitude of calcium oscillations was about 57 ± 19% (175 and 167 peaks, respectively) (**Fig. 9f**). The inhibitory effect of STIM1 clamping did not depend on the type of lipid anchor or potential lipid domain they may target. We performed the experiments using Lyn11-CIBN-GFP as a plasma membrane anchor and obtained similar results (**Supplemental Fig. 5a-b).** These results suggest that the reversibility of STIM1 translocation to the cortex is important for robust calcium oscillations (**Fig. 9g**). Strikingly, with both the CIBN-GFP-CAAX and Lyn11GFP-CIBN-GFP, we have observed in some cells where calcium oscillations still persisted after STIM1-Cry2 clamping but displayed reduced amplitudes (**Fig. 9h, Supplemental Fig. 5b)**, strongly suggesting the presence of an antagonistic role of STIM1 and cER translocation in oscillatory calcium responses (**Fig. 9i)**.

## Discussion

Calcium oscillations are the most well-known oscillatory reaction documented at the single cell level (*1, 11, 36*), but the kinetic mechanisms regulating the oscillations remain unclear. Dynamics of molecular machineries regulating SOCE under physiological conditions leading to calcium oscillations has been sparsely investigated. Compared to persistent store depletion that leads to stable, large-scale redistribution of STIM1 and Orai1/CRACM1, stimulation by antigen causes a more subtle yet complex pattern of STIM1 dynamics that are more subtle (*17, 37*). By characterizing STIM1 dynamics in single RBL-2H3 cell, we show that STIM1 displays two types of cyclic patterns, one of which is a travelling wave that is uncoupled from calcium oscillations. Previous studies support the model that each pulse of calcium oscillations stimulates STIM1 clustering and translocation to the cortex by depleting the internal stores and presumably after calcium refill. Our data suggest that in addition to the calcium-dependent mechanism, there is a second cortex-dependent oscillator that dictates STIM1’s cyclic translocation to the plasma membrane. The PI(4)P-dependent cycling of STIM1 is coupled with dynamic ER-PM contact formation and can work both independently of calcium oscillations (**Fig. 3b**) or be superimposed on calcium oscillations (**Fig. 3d**).

Previous studies on the cortical factors for STIM1 translocation have relied mostly on loss-of-function experiments and argue for a central role of PI(4,5)P2 in its mobilization. Acute or chronic loss of SOCE upon artificial and drastic reduction of PI(4,5)P2 would only indicate its contribution to SOCE but does not directly address which stage it may act or prove it as the rate-limiting step during event of activation. We conclude a critical role of PI(4)P in inducing STIM1 translocation based on three lines of evidences. Firstly, pulses of STIM1 translocation are synchronized with PI(4)P levels and precede the rise of PI(4,5)P2 levels. Secondly, STIM1 pattern differs from PI(4,5)P2 effectors, including FBP17, in their localizations and amplitude changes in oscillations. Thirdly, optogenetic manipulation that increases PI(4)P while decreases PI(4,5)P2 induces STIM1 translocation. The requirement for PI(4)P in STIM1 translocation is consistent with a number of studies based on pharmacological perturbations by inhibiting PI4K using high concentration of wortmannin or ly294002 (both are rather non-specific to PI4K) (*38*-*41*). Genetics manipulations showed STIM1 translocation to ER-PM contact sites upon thapsigargin treatment was impaired in PI4KIIIα knock-out cells (*21*). Because both PI(4)P and PI(4,5)P2 levels are likely coupled, the specific role of PI(4)P separate from that of PI(4,5)P2 remains difficult to pinpoint unambiguously. Our current study investigated their relationships under the more physiological conditions of calcium oscillations. More importantly, oscillations provided additional information on phase and amplitude responses which allowed us to tease apart the effects of PI(4)P and PI(4,5)P2.

PI(4)P oscillations have been observed at the trans-Golgi network (TGN) and was proposed to arise by PI4KIIα-dependent synthesis coupled with oxysterol-binding protein-dependent consumption (*42, 43*). The mechanisms underlying PI(4)P oscillations at the plasma membrane are not known. A key lipid enzyme for the synthesis of PI(4)P at the plasma membrane is PI4KIIIα (*21, 44, 45*) but it is unknown if it generates PI(4)P in a cyclical manner. On the other hand, PI(4)P can be degraded by the lipid phosphatase Sac1 which is localized to the ER, but may not be able to work in trans on substrates at the plasma membrane (*46*-*48*). Lipid exchange proteins such as ORP5 and ORP8 could contribute to PI(4)P metabolism by transporting it to the ER for Sac1-mediated degradation (*49, 50*). Whether these mechanisms are able to fulfil the much faster turnover rate in the plasma membrane PI(4)P oscillations (cycle time in tens of seconds compared to a few minutes at the TGN) requires further investigation. Using PI(4)P oscillations as a readout, one could potentially better understand the crosstalk between PI(4)P levels and ER-PM membrane contact sites formation.

Although our data does not suggest PI(4,5)P2 to be rate-limiting in the translocation of STIM1 to the cortex, they do not contradict previous data showing the importance of PI(4,5)P2 in STIM1 translocation (*20, 32, 40, 51*), or any downstream function of ER-PM contact sites formation in replenishing plasma membrane PI(4,5)P2 following receptor stimulation (*30, 52*). It is plausible that basal levels of PI(4,5)P2 are sufficient for STIM1 translocation under physiological conditions but lowering PI(4,5)P2 levels may limit the maximal level of STIM1 accumulation necessary for persistent store depletion. What our results emphasize is that the threshold for STIM1, as well as other cortical ER-PM contact proteins, could be lower than that of other PI(4,5)P2 and PI(3,4,5)P3 binding proteins. Even without the contribution of calcium-dependent factors, STIM1 translocation could proceed without an acute increase or even decrease in PI(4,5)P2 levels while translocation of other cytosolic proteins would require *de novo* PI(4,5)P2 synthesis to reach a higher threshold. Whether these different pools of PI(4,5)P2 reflect presence of PI(4,5)P2 nano-domains is less clear. Our data on the segregation of STIM1 and FBP17 is consistent with previous data showing separation of ORAI cluster from PI(4,5)P2 (*32*), and these data appear to suggest that STIM1 would occupy PI(4,5)P2-poor domain even if such domain exists. We therefore favor the idea that availability of lipids are regulated by chemical kinetics, which when coupled with rapid diffusion could function effectively as transient domains. Here, lifetime of lipids is dictated by the topology of the enzymatic networks, where the very transient presence of free lipids appear to be unavailable to effectors that are not in the immediate proximity.

Our study also suggests a role in the reversibility of ER-PM contact formation. Although it has not been previously appreciated, such reversibility is likely relevant for understanding the kinetics of calcium oscillations beyond the minimal machinery of SOCE. A number of optogenetics methods have been developed to artificially generate calcium responses by inducing the dimerization of STIM1 cytoplasmic domains (*53*-*59*). In none of these strategies was the reversibility of STIM1 specifically incorporated as part of the design, and yet calcium entry can be triggered by periodic light-induced dimerization of STIM1, suggesting that reversible association of STIM1 to the plasma membrane is not essential for generating a single wave of calcium entry. However, in these light-induced calcium responses, the kinetics of calcium entry is slow (rise time of 10 sec to minutes) and the total duration of the calcium pulses are minutes or longer (*60*). Under physiological conditions, oscillatory calcium spikes of the baseline spiking types have characteristic rise time between 1-3 sec, a total duration of about 7 sec and oscillatory periods between 10-30 sec (*11, 61*). Understanding the cooperativity needed for calcium entry in the oscillatory reaction likely requires considerations of the reversible and dynamic turnover of the ER-PM contact formation, which modulates the occurrence as well as kinetics and amplitude of calcium oscillations under physiological conditions.

## Materials and methods

### Plasmids and reagents

The following reagents were purchased from commercial sources: mouse monoclonal anti-dinitrophenol (DNP) IgE and thapsigargin were bought from Sigma-Aldrich (USA). DNP conjugated bovine serum albumin (DNP-BSA) was from Invitrogen (USA). Constructs of the following plasmids were kind gifts: iRFP-PH_PLCδ_, mCherry-PH_Grp1_, EGFP-C2-PH_FAPP1_, mCherry-CRY2-5-ptase_INPP5E_, from Pietro De Camilli (Yale University); Tks4-GFP from Begona Diaz (Sanford-Burnham Medical Research Institute); STIM1-RFP from David Allan Holowka (Cornell University); EGFP-Septin4 from Patrick Hogan (Harvard University School of Medicine); mEGFP-ER-5a from Michael Davidson (addgene #56455). GCaMP3 (#22692), R-GECO1 (#32444), mCherry-Sec61b (#90994), GFP-MAPPER (#117721), mRFP-wGBD (#26733), CIBN(deltaNLS)-pmGFP (shown as CIBN-GFP-CAAX in text)(#26867), Lyn11-CIBN-GFP (#79572), iRFP-P4M_SidM_ (#51470), iRFP (#31857) pmiRFP670-N1 (#79987) from Addgene. EGFP-C2-tandemPH_FAPP1_ was prepared by subcloning PH_FAPP1_ into the EGFP-C2-PH_FAPP1_ vector using restriction sites BglII and HindIII. miRFP670 was obtained by PCR from pmiRFP670-N1 using primer pairs CMV forward and CAAGCGGCCGCACTGCTCTCAAGCGCGGTGATCCGCG. The PCR product was then used to replace mCherry via Nhe1 and Not1 to obtain miRFP670-Cry2-ptase_INPP5E_. STIM1 was moved into piRFP using Xho1 and EcoR1 to obtain STIM1-iRFP. Cry2 was obtained by PCR from mCherry-Cry2-ptase_INPP5E_ using primer pair CAAAGTCGACAATGAAGATGGACAAAAAGACTATAG and CAAACCGCGGGGCTGCTGCTCCGATCATGATCTG. The PCR product was then inserted in between STIM1 and iRFP using Sal1 and Sac2. pmiRFP670-N1 was digested with Sac2 and Not1 and used to replace iRFP to obtain STIM1-Cry2-miRFP670. STIM1-Δpolybasic domain was obtained by PCR from STIM1-RFP using primer pair CMV forward and CAAAGGTACCCCTGGGCTGGAGTCTGTTTC. FBP17-EGFP and iRFP-N-WASP were previously described (*5, 31*).

### Cell culture and transfection

RBL-2H3 cells (ATCC) were maintained in monolayer culture in MEM (Invitrogen) supplemented with 20% fetal bovine serum (Sigma-Aldrich), and harvested with TrypLE Express (Invitrogen). For transient transfections, 2×10^6^ cells were collected and resuspend in 10μl R buffer (Invitrogen), 1μg DNA of each plasmid construct were added and electroporated using the Neon transfection system (Invitrogen) following the manufacturer’s instructions (1200 volts, 20-ms pulse width for 2 pulses). After transfection, cells were plated in 35mm glass bottom culture dishes (MatTek) and sensitized overnight with 0.4μg/mL anti-DNP IgE. Stable cell lines of FBP17-EGFP (*31*) or GCaMP3 were used for their respective imaging experiments and were generated by transfection of RBL-2H3 cells for 24 h, then selected in MEM with 0.5 mg/ml G418 (Invitrogen) and sorted for fluorescence. Cells were washed three times with Tyrodes solution (20mM HEPES-NaOH pH 7.4, 135mM NaCl, 5.0mM KCl, 1.8mM CaCl_2_, 1.0mM MgCl_2_ and 5.6mM glucose) before imaging. For antigen stimulation, DNP-BSA was diluted to a final concentration of 80ng/mL in the appropriate buffer and added to anti-DNP IgE sensitized cells. The time antigen is added is stated in the specific figure legends. For calcium store depletion, cells were stimulated with 2*μ*M or 10*μ*M thapsigargin.

### Perfusion experiments

GCaMP3 stable expression cells, or co-transfected with STIM1 construct, were cultured on No.1.5, 20mm diameter, round glass coverslips (Marienfeld-Superior) overnight. The coverslips are then transferred to a customized perfusion chamber to form the bottom of the chamber (Chamlide, Live Cell Instrument). A multi-valve perfusion control system (MPS-8, Live Cell Instrument) was used to switch rapidly between prewarmed, calcium-containing Tyrodes buffer or calcium-free buffer (20mM HEPES-NaOH pH 7.4, 135mM NaCl, 5.0mM KCl, 1.0mM MgCl_2_, 5.6mM glucose and 0.5 mM EGTA).

### Imaging

TIRFM was performed using a Nikon Ti inverted microscope with 3 laser lines (491nm, 561nm, 642nm) at 37°C. The microscope was equipped with an iLAS2 motorized TIRF illuminator (Roper Scientific). All images were acquired through Nikon objective (Apo TIRF 60X, N.A. 1.49 oil), a quad-bandpass filter (Di01-R405/488/561/635, Semrock) and a Prime 95B CMOS camera (Photometrics). The microscope, camera and illuminator were controlled by Metamorph 7.8 software (Universal Imaging). Cells were incubated in a stage-top heated incubator (Live Cell Instrument) set at 37°C and were typically imaged at 2sec per frame for live imaging movies.

### Optogenetics and fluorescence recovery after photobleaching (FRAP)

For optogenetics experiments, we followed previously published methods(*31*). Membrane-anchored probe CIBN-GFP-CAAX or Lyn11-CIBN-GFP was used to recruit its cytosolic binding partner Cry2 fused with target proteins upon 491nm light activation. For all the constructs, mCherry-Cry2-5-ptase_INPP5E_, miRFP670-Cry2-ptase_INPP5E_ and STIM1-Cry2-train of pulses with intervals varying between 20sec to 1min was used to achieve instant or sustained Cry2 recruitment, respectively. We noticed that the calcium sensor GECO1 undergoes reversible photoactivation when stimulated by 491nm laser at powers higher than 0.2mW, therefore we restricted 491nm laser power to 0.1mW for optogenetics to prevent photoactivation of GECO1 for Fig. 7d-e, Fig. 9d,e,h and Supplemental Fig. 5. For optogenetics on calcium oscillations, we optimized the experimental conditions in order to obtain uniform and robust calcium oscillations. Only RBL-2H3 cells with passage number less than eight were used. We observed that transient transfection of three plasmids together [CIBN-GFP-CAAX (1ug), GECO1 (1ug) and STIM1-Cry2-miRFP670 (3ug)] or [Lyn11-CIBN-GFP (1ug), GECO1 (1ug) and STIM1-Cry2-miRFP670 (3ug)] induce more cell death compared to transfection of two plasmids. To obtain higher cell number for imaging, 2 to 3×10^6^ cells were seeded in one 35mm glass bottom culture dishes (MatTek) after transient transfection. Cells were stimulated with antigen in calcium-free buffer and exchange to calcium-containing buffer as shown in Fig. 1. For fluorescence recovery after photobleaching (FRAP) experiment (Fig. 4e), cells were imaged at 2sec per frame and photobleaching was performed using maximum power (10mW) of 491 nm laser for 1sec at the region of interest.

### Imaging analysis and statistics

Post-acquisition imaging analysis was performed using Fiji and Matlab (Mathworks). A “reslice” tool and “median” projection filter in Fiji were used to generate kymographs. A “montage” tool was used to generate montages of selected region of interest (ROIs). Pseudocolor images in Fig. 2a and 4e were generated using “Thermal”, and montage in Fig. 4d and kymograph in Fig. 7e and 9d using “Gem” lookup table in Fiji. To identify the frequency in Matlab was used. FFT trace showing a major peak with three times higher amplitude than the second highest peak is considered as oscillations. To determine the phase shift between different probes in Fig. 5, 6 and Supplementary Fig. 3, intensity profiles in a region of interest (15×15 pixels) were smoothed (smoothspan ranging from 5sec to 8sec). Multiple cycles were aligned based on the first peak of FBP17 (Fig. 5), N-WASP (Supplemental Fig. 3), PH_Fapp1_ (Fig. 6a), PH_PLC_ (Fig. 6b), PH_Grp1_ (Fig., 6c), Tks4 (Fig. 6d) intensity, and time delay between two probes was quantified by cross-correlation analysis in Matlab. For FRAP quantification in Fig. 4e, the fluorescence intensity of the region of interest after bleaching was normalized to the mean intensity before bleaching. For comparison of fluorescence increase, the fluorescence intensity were quantified with a region of interest (30×30 pixels) after background subtracted, percentage of fluorescence intensity changes is quantified by (Ft_cell_-F0_cell_)/(F0_cell_-F0_background_) in Fig. 9b, fold change is quantified by (Ft_cell_-F0_background_)/(F0_cell_-F0_background_) for Fig. 4a-b, amplitude is quantified by Ft_cell_-F0_cell_ in Fig. 7b-d, 9c and Supplementary Fig. 2a-b. For comparison of the same sample before and after the perturbation in Fig. 7b-d and 9c, paired two-tailed Student *t*-test was used. For comparison between two groups in Fig. 7e-f, 9f and Supplemental Fig. 2, unpaired two-tailed Student *t*-test was used, average data is shown as mean±s.d.

## Data availability

All data supporting the findings of this study are available from the corresponding author upon reasonable request.

## Code availability

Computer code used for fast Fourier transform and phase correlation are available upon request.

## Acknowledgement

We would like to thank Qiang Yuan and Zhifu Wang for assistance with video editing, Su Guo for assistance with molecular biology and Emma Feng for assistance with imaging. This work was supported by the Singapore Ministry of Education Academic Research Fund Tier 2 (M. Wu), the Singapore Ministry of Health National Medical Research Council Open Fund Individual Research Grant (M. Wu, NMRC/OFIRG/0038/2017), Yale University startup grant (M. Wu).

## Supplementary Materials

**Supplemental Fig. 1.**
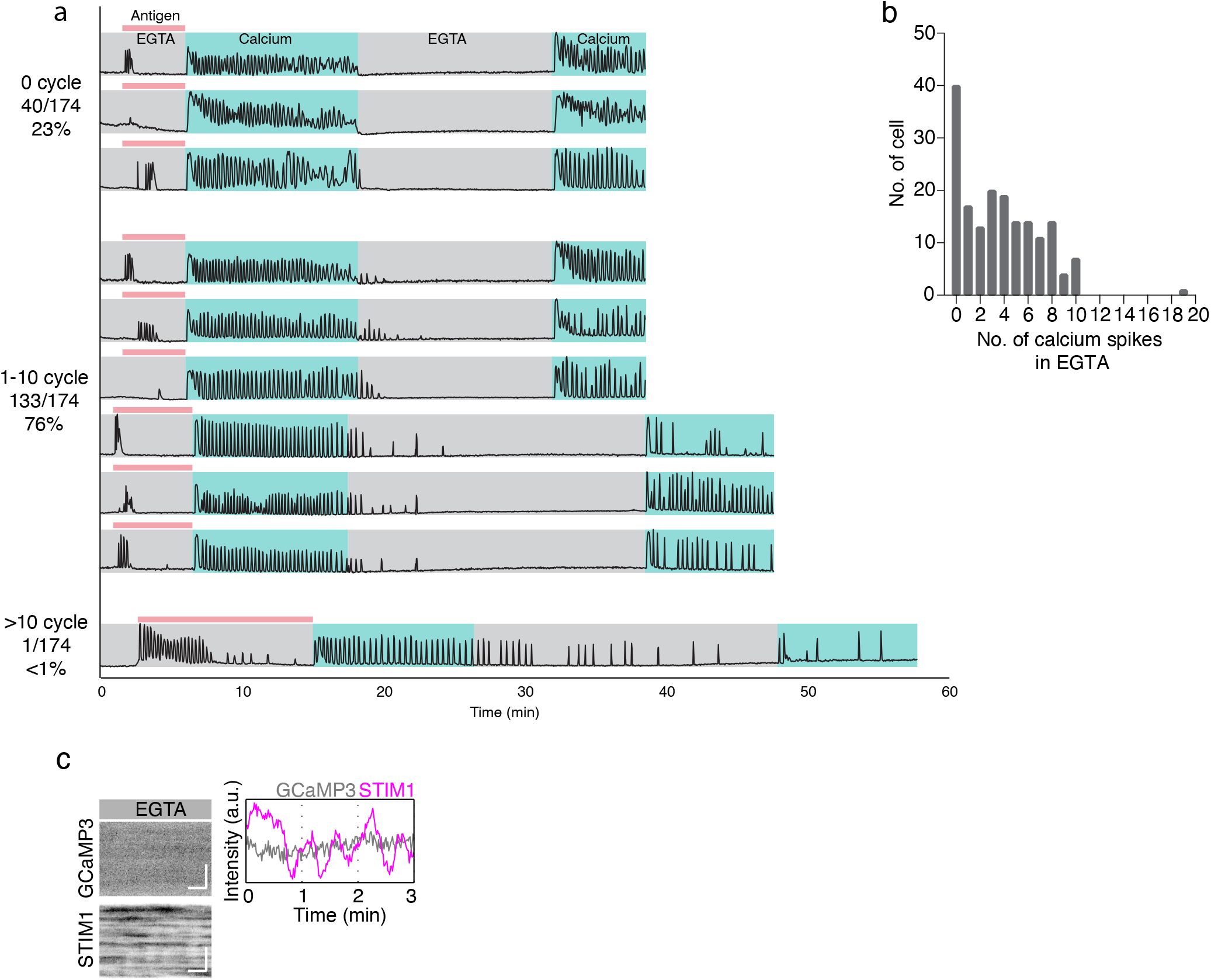
Extracellular calcium is required for sustained calcium oscillations upon antigen stimulation. (a) Intensity profiles and histogram of cells stably expressing GCaMP3 perfused with alternating solutions containing calcium, EGTA or EGTA with antigen. The proportion of cells with the indicated numbers of calcium spikes after change from calcium-containing to EGTA buffer is stated on the left. Images are acquired 2sec/frame (n=189 cells from 5 experiments). (b) Histogram of cells with various numbers of calcium spikes in EGTA after antigen stimulated calcium oscillations (174 cells from 5 experiments). (c) Kymographs and intensity profile of GCaMP3 and STIM1-RFP in EGTA buffer 10min after antigen stimulation (3 cells from 3 experiments). Scale bar=30 sec (horizontal bar), 5 µm (vertical bar).

**Supplemental Fig. 2.**
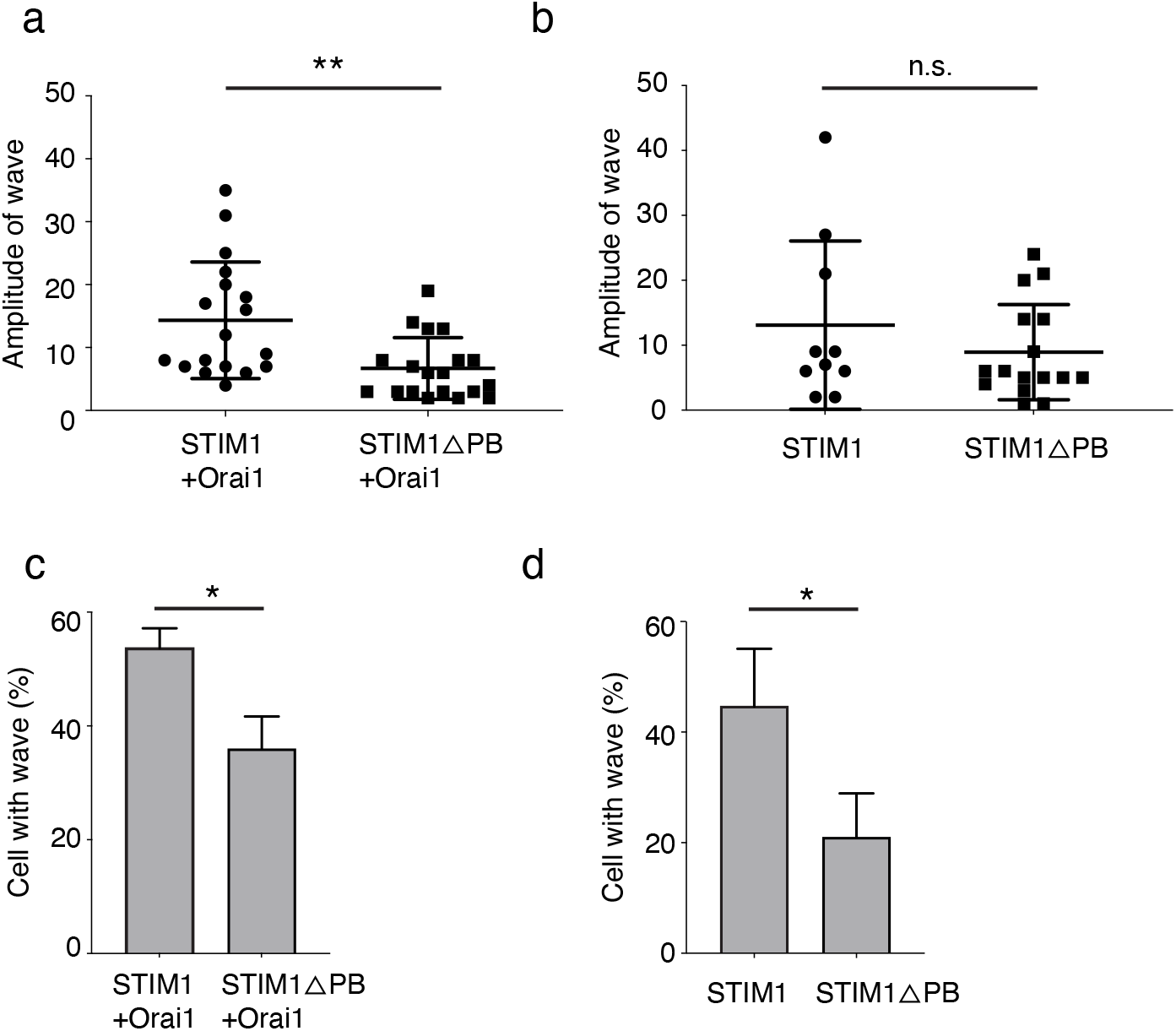
STIM1 translocation is reduced with Δpolybasic domain. (a-b) Quantification of the amplitude of cell displaying wave upon antigen stimulation in cell over expressing STIM-RFP/STIM1-Δpolybasic domain with Orai1 (a) (n=18, 19 cells from 3 experiments, respectively) or alone (b) (n=10, 15 cells from 3 experiments, respectively). (c-d) Quantification of percentage of cells displaying waves upon antigen stimulation in cell over-expressing STIM1-RFP/STIM1-Δpolybasic domain with Orai (c) (n=41, 52 cells from 3 experiments, respectively) or alone (d) (n=46, 72 cells from 3 experiments, respectively). Amplitude is quantified by Ft_cell_-F0_cell_. Error bar: s.d.; n.s., nonsignificant, * *P*<0.05, ** *P*<0.01, *t*-test. Scale bar=30 sec (horizontal bar), 5 µm (vertical bar).

**Supplemental Fig. 3.**
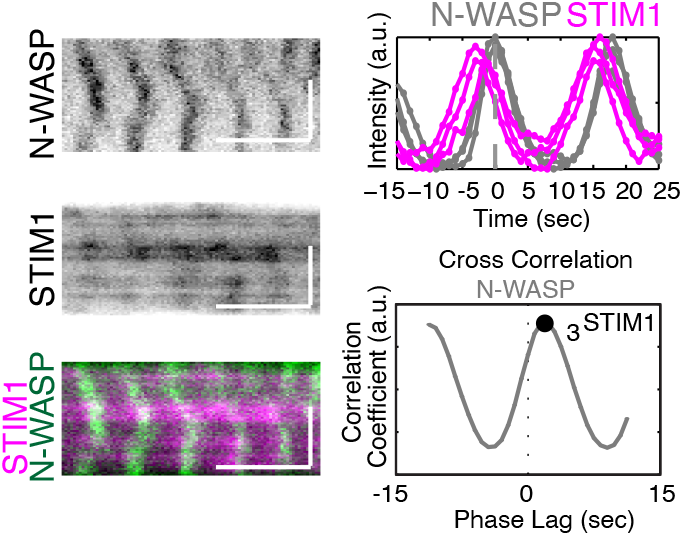
Coupling of STIM1 with cortical protein in wave dynamics. Kymographs, intensity profile and cross-correlation analysis of a cell displaying iRFP-N-WASP and STIM-RFP waves, 10min after antigen (n= 7 cells from 3 experiments). Scale bar=30 sec (horizontal bar), 5 µm (vertical bar).

**Supplemental Fig. 4.**
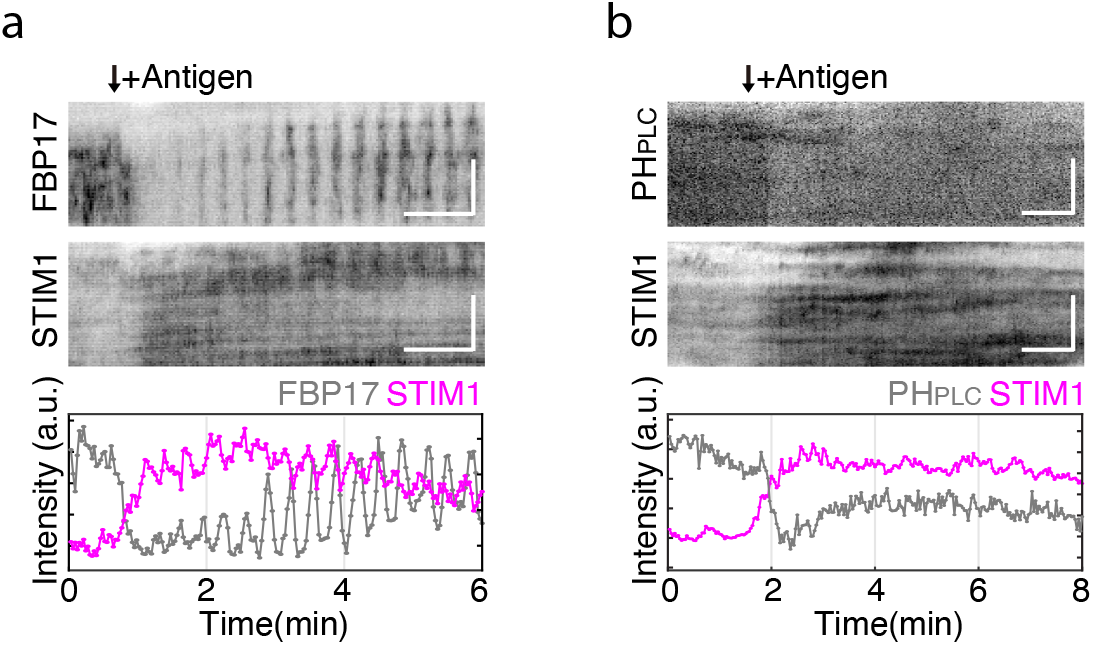
Dynamics of STIM1 with FBP17 and PI(4,5)P2 in the initial phase of antigen stimulation. (a) Representative kymographs and intensity profiles of a cell stably expressing FBP17-EGFP and co-transfected with STIM1-RFP (n=8 cells from 3 experiments). (b) Representative kymographs and intensity profiles or iRFP-PH_PLCδ_ and STIM1-RFP upon antigen stimulation (n=22 cells from 5 experiments). Scale bar=1 min (horizontal bar), 5 µm (vertical bar).

**Supplemental Fig. 5.**
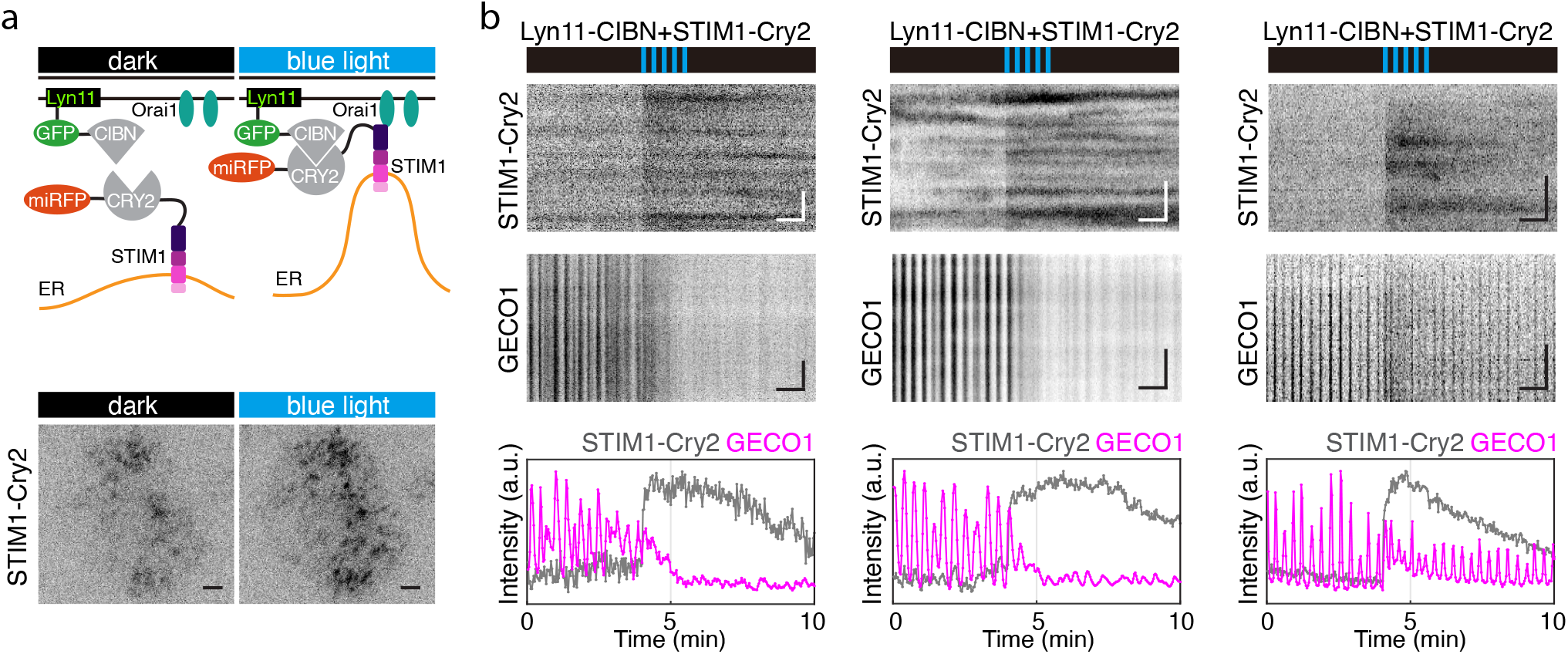
Optogenetic translocation of STIM1 by Lyn11-CIBN-GFP regulates calcium oscillations. (a) *Top*: Schematic of light-induced plasma membrane translocation of STIM1-Cry2-miRFP670 by Lyn11-CIBN-GFP. *Bottom*: Representative TIRF images of a cell before and after blue light exposure without antigen stimulation (20 cells from 4 experiments). (b) Kymographs and intensity profile of STIM1-Cry2-miRFP670 and GECO1 during exposure to blue light pulses, movies were acquired 10 to 30min after antigen stimulation. Cell showing almost abolished or decreased calcium oscillations when STIM1-Cry2-miRFP670 is recruited upon blue light pulses (15 cells from 4 experiments). For TIRF images, scale bar=5 µm; for kymographs, scale bar=1min (horizontal bar), 5 µm (vertical bar).

**Supplemental video** 1. Calcium oscillations are dependent on extracellular calcium. TIRF movie of cell stably expressing GCaMP3 pre-incubated in EGTA containing buffer, stimulated with antigen, and change to calcium containing buffer. The video was acquired at 2 sec interval and played at 30x real time. Scale bar: 10 µm.

**Supplemental video** 2. Cyclic STIM1 translocation with or without corresponding calcium signals. Two-color TIRF movie of cell expressing GCaMP3 and STIM1-RFP showing (A) cyclic SITM1 translocation with calcium oscillations, GCaMP3 in grayscale or pseudocolor, and STIM1-RFP in pseudocolor; (B) cyclic STIM1 translocation with irregular calcium spikes; (C) cyclic STIM1 translocation without calcium spikes. Videos were acquired 10-40 min after antigen stimulation at 2 sec interval and played at 30x real time. Intensity profiles of selected regions of interest are shown accordingly. Scale bar: 10 µm.

**Supplemental video** 3. Two-color TIRF movie of cell expressing STIM1-RFP and mEGFP-ER-5a showing coupled traveling waves. The video was acquired 10 min after antigen stimulation at 2 sec interval and played at 30x real time. Scale bar: 10 µm.

**Supplemental video** 4. Two-color TIRF movie of cell expressing FBP17-EGFP and STIM1-RFP showing traveling waves. The video was acquired 15 min after antigen stimulation at 2 sec interval and played at 30x real time. Scale bar: 10 µm.

**Supplemental video** 5. Two-color TIRF movie of cell expressing CIBN-GFP (not imaged), miRFP670-Cry2-ptase_INPP5E_, and GECO1 showing inhibition of calcium oscillations upon optogenetic recruitment of miRFP670-Cry2-ptase_INPP5E_ to plasma membrane. The movie was acquired 20 min after antigen stimulation at 2sec interval and played at 30x real time. Scale bar: 10 µm.

**Supplemental video** 6. Two-color TIRF movie of cell expressing CIBN-GFP (not imaged), STIM1-Cry2-miRFP670, and STIM1-RFP showing optogenetic recruitment of STIM1-Cry2-miRFP670 cause coupled STIM1 translocation. The movie was acquired without antigen stimulation at 2sec interval and played at 30x real time. Scale bar: 10 µm.

**Supplemental video** 7. TIRF movie of cell expressing STIM1-Cry2-miRFP670 showing the plasma membrane translocation upon 10*μ*M thapsigargin. The movie was without antigen stimulation at 2sec interval and played at 30x real time. Scale bar: 10 µm.

